# Boundary updating as a source of history effect on decision uncertainty

**DOI:** 10.1101/2023.02.28.530543

**Authors:** Heeseung Lee, Sang-Hun Lee

## Abstract

When sorting a sequence of stimuli into binary classes, current choices are often negatively correlated with recent stimulus history. This phenomenon—dubbed the repulsive bias—can be explained by boundary updating, a process of shifting the class boundary to previous stimuli. This explanation implies that recent stimulus history can also influence “decision uncertainty,” the probability of making incorrect decisions, since it depends on the location of the boundary. However, there have been no previous efforts to elucidate the impact of previous stimulus history on decision uncertainty. Here, from the boundary-updating process that accounts for the repulsive bias, we derived a prediction that decision uncertainty increases as current choices become more congruent with previous stimuli. We confirmed this prediction in behavioral, physiological, and neural correlates of decision uncertainty. Our work demonstrates that boundary updating offers a principled account of how previous stimulus history concurrently relates to choice bias and decision uncertainty.

## INTRODUCTION

We often characterize a stimulus for a particular dimension by assigning a binary class, such as ‘small or large’ for size, ‘slow or fast’ for speed, and ‘light or heavy’ for weight. This seemingly natural and basic linguistic act entails a cognitive computation called ‘binary classification,’ where we decide which side of the quantity distribution the stimulus falls on, with respect to the dimension of our interest^1-4^. Thus, successful binary classification requires not just accurately perceiving the stimulus quantity but also knowing the class boundary that divides the quantity distribution^1^.

However, in everyday situations, the boundary is seldom presented explicitly at the moment of classification and, thus, needs to be learned and internally formed by observers themselves^5-12^.

Importantly, it has been suggested that this internal boundary tends to continually shift towards recent stimuli^7-10,12^ (Figure 1A). This “boundary shift towards recent stimuli” has accounted for a well-known history effect on choice called ‘repulsive bias,’ where the current choice is biased away from previous stimuli on a trial-to-trial basis^8-10,12,13^ (Figure 1A,B). Recently, we formalized this boundary updating in the framework of Bayesian decision theory—called ‘Bayesian Model of Boundary Updating (BMBU) — and demonstrated that BMBU readily accounts for the repulsive bias not just in behavioral choice but also in brain activity patterns^12^.

**Figure 1.**
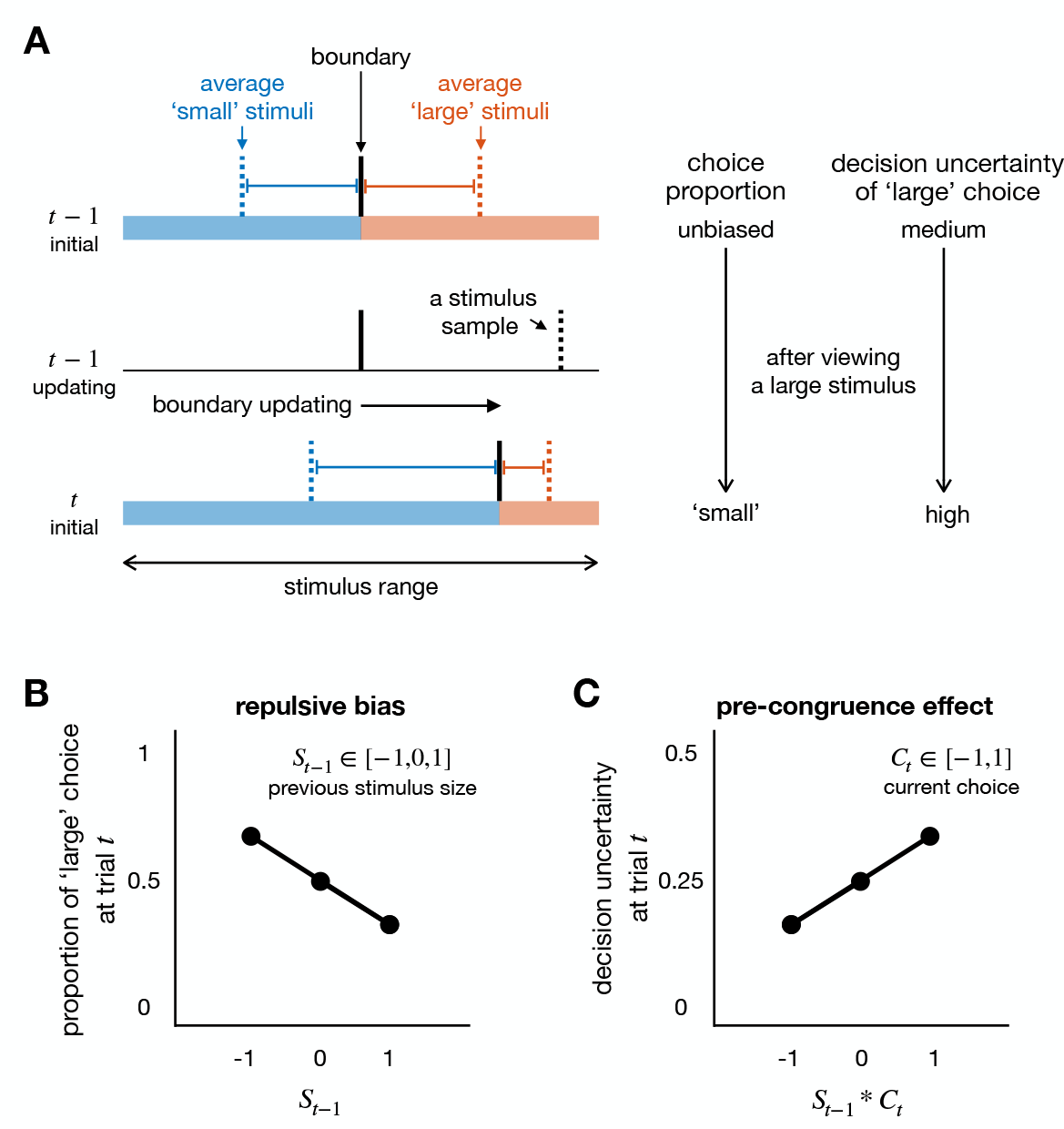
Boundary updating, as a common source of bias in choice and decision uncertainty during binary classification. (A) Schematic illustration of how boundary updating leads to the repulsive bias in choice and the pre-congruence effect on decision uncertainty. When the class boundary (solid vertical black bars) is unbiased, the stimulus sizes that lead to ‘large’ choices and those to ‘small’ choices are equal in proportion (as indicated by a large blue rectangle and a small red rectangle) and in distance (as indicated by a long blue horizontal line and a short red horizontal line) to the boundary (top row, left). Thus, both choice and decision uncertainty are unbiased (top row, right). According to the hypothesis of boundary updating, the boundary shifts toward the large side after encountering a large stimulus (middle row, left). As a result, the stimulus sizes that lead to ‘large’ choices and those to ‘small’ choices are no longer equal both in proportion and in distance to the boundary (bottom low, left). The imbalance in proportion leads to more ‘small’ choices in the following trial (the repulsive bias; bottom row, right), while the imbalance in distance to the boundary leads to an increase in decision uncertainty for the ‘large’ choice in the following trial (the pre-congruence effect; bottom row, right). Horizontal rectangles demarcate the stimulus ranges that are smaller (blue) or larger (red) than the boundary. The dotted blue and red vertical bars demarcate the average stimulus sizes smaller and larger than the boundary, respectively. (B) Repulsive bias in choice proportion. The repulsive bias can be signified by the negative correlation between the choice in the current trial and the stimulus in the previous trial. *S*_*t*-1_ indicates the stimulus size at trial *t* − 1 (the immediately preceding trial), where small, medium, and large sizes are denoted by −1, 0, and 1, respectively. (C) Pre-congruence effect on decision uncertainty. The pre-congruence effect can be signified by the positive correlation between the decision uncertainty at trial *t* (the current trial) and the congruence of the choices between the previous and current trials. *C*_*t*_ indicates the choice at trial *t*, where ‘small’ and ‘large’ choices are denoted by −1 and 1, respectively.

Here, we note that the same boundary updating process that induces the repulsive bias in ‘choice’ should also confer another history effect on ‘decision uncertainty.’ Decision uncertainty is the probability that the current choice is incorrect, and this probability increases as the stimulus approaches nearer the boundary in binary classification tasks^11,14-19^. Suppose the boundary was initially neutral (Figure 1A, top), and a large stimulus was presented in the last trial (Figure 1A, middle). The boundary would then shift towards the large side according to the boundary updating hypothesis^7-10,12^, bringing itself nearer to the stimuli associated with the ‘large’ choice but farther from those associated with the ‘small’ choice in the following trials (Figure 1A, bottom). Consequently, having previously viewed large stimuli, observers’ ‘large’ choices are likely to be accompanied by high levels of uncertainty, whereas their ‘small’ choices by low levels of uncertainty. Generally put, the more congruent the current choice is with previous stimuli, the more uncertain it is (Figure 1C). We shall call this boundary-induced history effect on decision uncertainty *the pre-congruence effect*.

It is surprising that this seemingly straightforward, pre-congruence effect has not been recognized theoretically nor demonstrated empirically since decision uncertainty plays a fundamental role in diverse adaptive behaviors^20-22^. Decision uncertainty has been considered a crucial cognitive variable in adaptive human behavior, such as volatility monitoring^20^, learning from errors^21^, and executive control^22^. It should be noted that decision uncertainty differs from sensory uncertainty, which is influenced solely by the noise present in stimuli^23^, whereas decision uncertainty can be modulated by the distance between the stimulus and the boundary even if sensory uncertainty is constant. Furthermore, decision uncertainty has wide-ranging positive associations with response time^11,16-18,24^, pupillary responses^18,25-34^, and neural activities in various brain regions^15,35-44^.

To find empirical evidence for the pre-congruence effect, we probed the response time (RT), pupil size, and functional magnetic resonance imaging (fMRI) signals of human observers during binary classification tasks with diverse experimental conditions. Initially, we confirmed that the current datasets display the history effect on choice (i.e., the repulsive bias) as predicted by BMBU. Then, by simulating BMBU, we quantitatively derived a set of implications of boundary updating on decision uncertainty, including the pre-congruence effect. Next, we confirmed these model implications by conducting multiple regression analyses on the RTs, the cortical activity in the dorsal anterior cingulate cortex (dACC), and the pupil-size measurements. The pre-congruence effect was significant in all datasets analyzed, and it remained robust despite changes in sensory modality, stimulus granularity, inter-trial interval, and trial-to-trial feedback. The magnitude of the pre-congruence effect was substantial, ranging from 10% to 45% of the effect of the current stimulus. Furthermore, the pre-congruence effect remained unaffected in the presence of previously known history effects on decision uncertainty. These results point to ‘boundary updating towards recent stimuli’ as a common source of two types of history effects: the repulsive bias, a well-known history effect on choice, and the pre-congruence effect, a previously unrecognized history effect on decision uncertainty.

## RESULTS

### History effects of previous stimuli on the current choice

Forty-one human individuals participated in our study and performed a binary classification task. They viewed a series of rings that appeared in a pseudo-randomized order^45^ while classifying the size of each ring as ‘small’ or ‘large’ under moderate time pressure (Figure 2A). As previously reported^7-10,12^, the participants showed the repulsive bias by making the ‘small (large)’ choice more often after seeing the large (small) stimulus than after seeing the small (large) stimulus in the previous trial (empty circles in Figure 2B). The repulsive bias was supported by the significant negative regression of the current choice *C*_*t*_ onto the previous stimulus 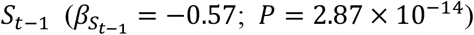 in a multiple logistic regression model where *C*_*t*_ was simultaneously regressed onto the current stimulus 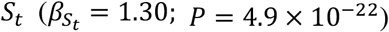 and the previous choice 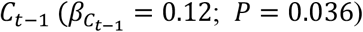. The repulsive bias was not constrained to the immediately preceding stimulus but extended to the stimulus at the lag of three trials (Figure 2C).

**Figure 2.**
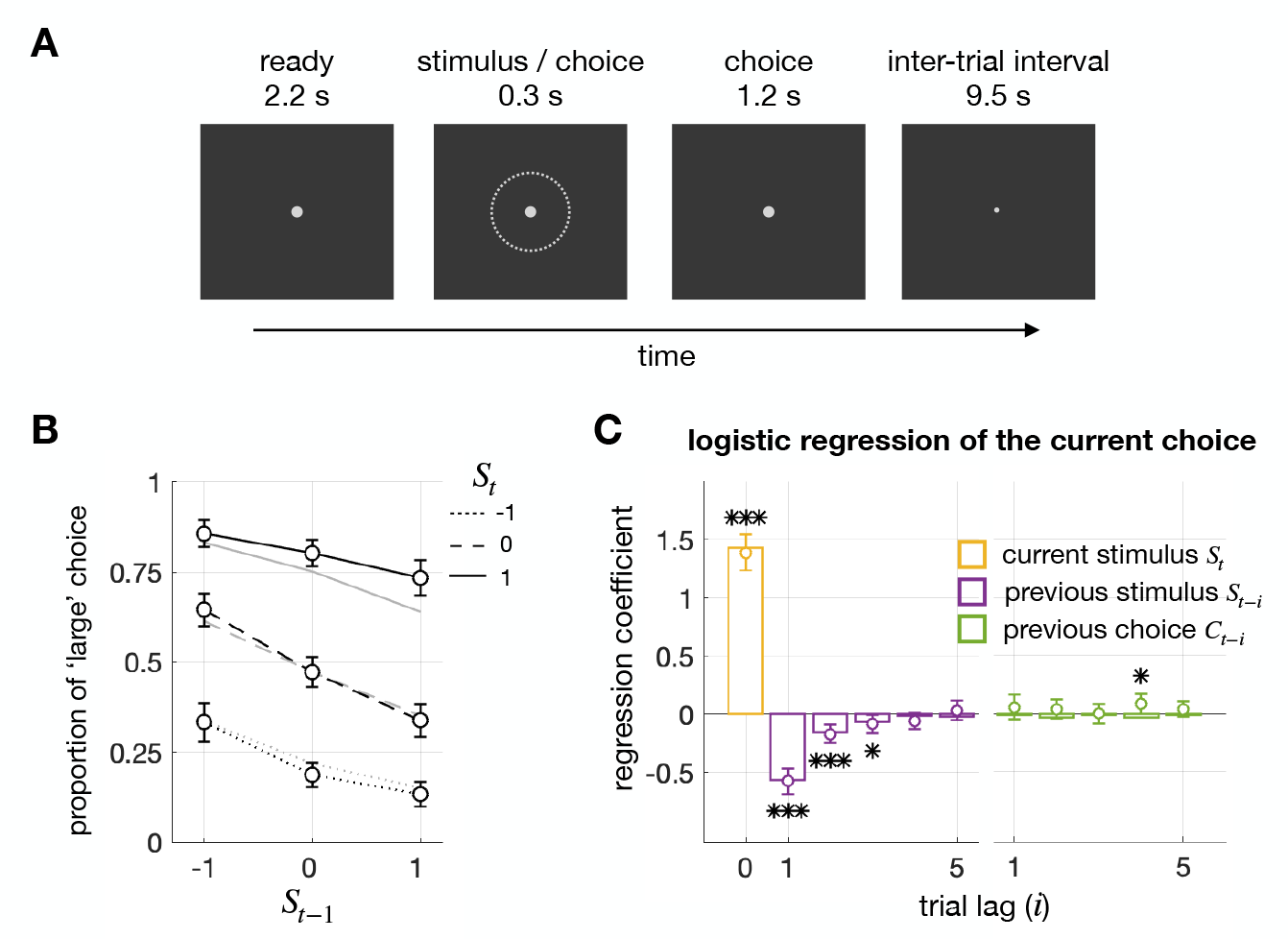
Task structure and repulsive bias. (A) Task structure. On each trial, the size of a ring was classified as ‘small’ or ‘large’ within 1.5 sec after the stimulus onset. (B) Repulsive bias in psychometric curves. The proportions of ‘large’ choices conditioned on current stimuli (*S*_*t*_) are plotted against previous stimuli (*S*_*t*-1_), in black lines for human observers and in gray lines for the Bayesian model for boundary updating (BMBU). The negative slopes of the curves signify the repulsive bias. (C) Repulsive bias in logistic regressions. The coefficients in the multiple logistic regression of current choices (*C*_*t*_) onto current and previous stimuli (*S*_*t*-*i*_) and previous choices (*C*_*t*-*i*_) are plotted in circles and bars for humans and BMBU, respectively. The significant negative coefficients of previous stimuli, not previous choices, signify the repulsive bias. Here and thereafter, error bars represent the 95% confidence interval of the means, and asterisks indicate *P* values of two-sided Student’s *t*-test: ∗< 0.05, ∗∗< 0.01, ∗∗∗< 0.001.

Previously, our group demonstrated that the repulsive bias can be explained by the continual updating of the class boundary toward previous stimuli^12^. In doing so, we formalized such boundary updating with a Bayesian model, called the ‘Bayesian model of boundary updating (BMBU),’ and showed that BMBU quantitatively reproduces the history effects on choice with an exquisite level of finesse (, as indicated by the gray lines in Figure 2B and the bars in Figure 2C).

In the current study, we will demonstrate that boundary updating can also explain the history effects on decision uncertainty. To start, we will provide a brief overview of BMBU (see STAR methods) and describe how boundary updating relates to decision uncertainty.

### Bayesian model of boundary updating

BUBU critically assumes that Bayesian observers have learned that their memory of stimuli becomes noisier as trials elapse^46,47^. Specifically, the learned generative model (Figure 3A) incorporates this memory decay by increasing the width of the distribution of noisy memory recalls 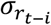 as a function of the number of elapsed trials *i* when the memory of the *i*-th previous stimulus *r*_*t-i*_ is recalled in the current trial *t*. The process of memory decay is approximated by a power function with a rate parameter *k* (*k* > 0):

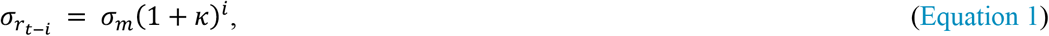

where σ_*m*_ is the width of the distribution of sensory measurements of the current stimulus *m*_*t*_.

**Figure 3.**
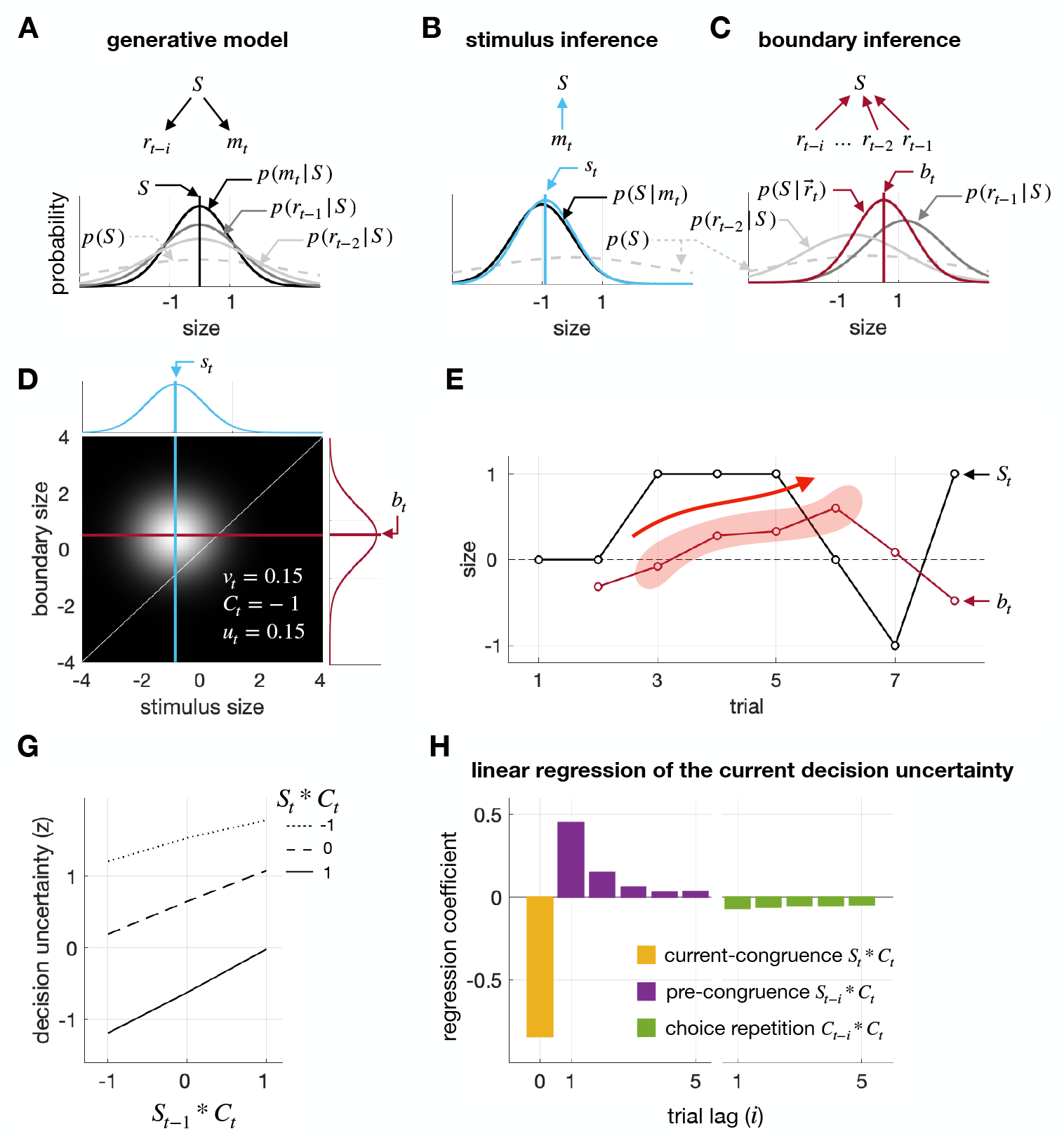
Bayesian model of boundary updating and its simulation of pre-congruence effect. (A) Generative model. BMBU posits that the sensory measurement of the current stimulus (*m*_*t*_) and the memory evidence of the previous stimulus in the *i*-th preceding trial (*r*_*t*-*i*_) are both normally distributed, as denoted by the solid curves *P*(*m*_*t*_|*S*_*t*_) and *P*(*r*_*t*-*i*_|*S*_*t*-*i*_), respectively. For illustrative purposes, the stimulus sizes are set at a single value (*S*_*t*_ = *S*_*t*-*i*_ = *S*). To account for the decay in memory precision, BMBU posits that the distribution of memory measurements *P*(*r*_*t*-*i*_|*S*_*t*-*i*_) becomes broad as trials elapse. Additionally, BMBU posits that observers have prior knowledge of the distribution of physical stimulus size, as denoted by the dashed curve *P*(*S*). (B) Inference of stimulus size. BMBU infers the current stimulus size by constructing the posterior distributions of stimulus size based on the current sensory measurement (*P*(*S*|*m*_*t*_)). The estimate of the stimulus *s*_*t*_ is the mean of the posterior distribution, as indicated by the blue vertical line. (C) Inference of boundary location. BMBU infers the boundary by constructing the posterior distribution of stimulus size based on the memory measurements of the previous stimuli 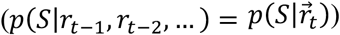. The estimate of boundary *b*_*t*_ is the mean of the posterior distribution, as indicated by the red vertical line. For illustrative purposes, only the memory measurements of the stimuli within trial lags of two (*r*_*t*-1_ and *r*_*t*-2_) are considered. Note that the boundary estimate (*b*_*t*_) is more biased toward the memory measurement of the more recent stimulus (*r*_*t*-1_) than the farther one (*r*_*t*-2_), which explains why the attraction of the boundary toward previous stimuli diminishes as trials elapse. (D) Binary classification with decision uncertainty. The posterior distributions of stimulus size and class boundary are projected onto a bivariate space, where the image intensity corresponds to the probability density of the product of the two posterior distributions 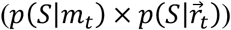. BMBU classifies the current stimulus as ‘large’ if the probability that the inferred stimulus is larger than the boundary (the decision variable, *v*_*t*_) is greater than 0.5, and as ‘small’ otherwise. Decision uncertainty (*u*_*t*_) is quantified as the probability that the current choice (*C*_*t*_) is incorrect and increases as the stimulus gets nearer the boundary. In this particular example, *v*_*t*_ is 0.15, *C*_*t*_ is ‘small’ (−1), and *u*_*t*_ is 0.15. (E) An example trajectory of the boundary in a representative subject 04. The black and red lines track the stimulus size and the boundary inferred by BMBU, respectively, over eight trials. The shaded red area highlights a gradual shift of the boundary towards the large side due to a streak of large stimuli. (G) Model simulation of pre-congruence and current-congruence effects in psychometric curves. The simulated decision uncertainty increases with the congruence between the current choice and the previous stimuli (*S*_*t*-1_ ∗ *C*_*t*_) but decreases with the congruence between the current choice and the current stimulus (*S*_*t*_ ∗ *C*_*t*_), the former and latter signifying the pre-congruence and current-congruence effects on decision uncertainty, respectively. (H) Model simulation of pre-congruence and current-congruence effects in linear regressions. The vertical bars represent the coefficients of the multiple linear regression of the simulated decision uncertainty onto the congruences of current choices with the current stimulus (*S*_*t*_ ∗ *C*_*t*_), the previous stimuli (*S*_*t*-*i*_ ∗ *C*_*t*_), and the previous choices (*C*_*t*-*i*_ ∗ *C*_*t*_). The pronounced positive coefficients of the congruence between the current choice and recent stimuli (purple bars) signify the pre-congruence effect. See also Figure S1.

With this generative model, the Bayesian observer can form two probabilistic beliefs, one regarding the current stimulus and the other regarding the current boundary, by inversely propagating the current sensory measurement *m*_*t*_ (Figure 3B) and the memory recalls of the previous stimuli *r*_*t-i*_ (Figure 3C), respectively, over the generative model. Note that the belief about the boundary can be formed in that way since the boundary for binary classification—a value that splits the stimulus distribution into even halves—is defined as the average size of the previous stimuli, which can be inferred from the noisy memory recalls of the observer. Here, the mean of this belief *b*_*t*_ is determined as the weighted average of recent stimulus memories *r*_*t-i*_ and a long-term stimulus memory *μ*_0_:

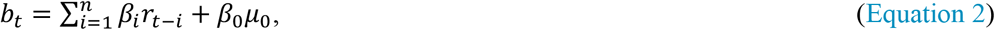

where the weights are determined according to the generative model, as follows: 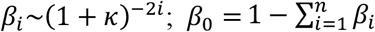. Consequently, past stimuli all attract *b*_*t*_ toward themselves, but with varying amounts: the attraction is maximal for the most recent one and decreases as trials elapse because *β*_1_ > *β*_2_ > … > *β*_*n*_.

Based on these beliefs about the current stimulus and boundary, the Bayesian observer computes the probability that the stimulus is larger than the boundary (the below-diagonal fraction of the bivariate distribution in Figure 3D), which is called decision variable. The decision variable *v*_*t*_ determines the current choice *C*_*t*_: *C*_*t*_ is ‘large’ (*C*_*t*_ = 1) when *v*_*t*_ is greater than 0.5 and ‘small’ (*C*_*t*_ = −1) otherwise.

Importantly, the decision uncertainty *u*_*t*_ can also be computed by assessing the probability that the choice is incorrect from the same beliefs (the below-diagonal fraction of the bivariate distribution in Figure 3D), which is the reverse of confidence (*f*_*t*_), the probability that the choice is correct (*u*_*t*_ = 1 – *f*_*t*_)^48,49^. Thus, according to BMBU, the decision uncertainty is inversely proportionate to the distance between the stimulus and boundary estimates.

### The attraction of the boundary to recent stimuli implies the repulsive bias for choice behavior and the pre-congruence effect for decision uncertainty

The way that the boundary is updated in BMBU implies that recent stimuli attract the boundary toward themselves on a trial-to-trial basis. To illustrate the trial-to-trial latent states of the inferred boundary, we fitted the parameters of BMBU to a single participant’s choice data and simulated the boundary estimates using the best-fit parameters. A snapshot example of the resulting simulation is shown in Figure 3E, where the implication of BMBU can readily be appreciated by continuous and immediate shifts of the simulated boundary estimates toward the recent stimuli.

This boundary attraction toward recent stimuli, in turn, has two ramifications for decision behavior in the context of history effects, one regarding ‘choice’ and the other regarding ‘decision uncertainty.’ Its ramification for choice behavior is the repulsive bias: the boundary being attracted to previous stimuli causes the current choice to be biased toward the side opposite to the previous stimulus, as intuited early on (Figure 1A,B) and shown previously by our group^12^.

The other—previously unidentified—ramification of the boundary attraction toward recent stimuli for decision uncertainty is the pre-congruence effect: the greater the congruency between the previous stimuli and the current choice (*S*_*t-i*_ ∗ *C*_*t*_) is, the greater the current level of decision uncertainty (*u*_*t*_) is (Figure 3G), as we intuited early on (Figure 1A,C). Since the contribution of the previous stimulus to boundary updating diminishes as trials elapse due to memory decay (Figure 3A), BMBU predicts that the pre-congruence effect accordingly decreases as trials further lag (the purple bars in Figure 3H), similarly to the repulsive bias (the purple bars in Figure 2C).

It should be noted that besides the previously unidentified pre-congruence effect, BMBU readily captures the well-established link of decision uncertainty with the congruency between the current stimulus and the current choice (*S*_*t*_ ∗ *C*_*t*_). This phenomenon, where decision uncertainty decreases as the current stimulus becomes increasingly congruent with the current choice^14,18,48^, will be referred to as the ‘current-congruence effect’ to distinguish it from the pre-congruence effect. BMBU predicts the current-congruence effect (as indicated by the progressive descent of the lines as a function of *S*_*t*_ ∗ *C*_*t*_ in Figure 3G and the yellow bar in Figure 3H) because the more congruent the current stimulus is with the current choice, the farther away it is from the current boundary, and thus the lower the decision uncertainty.

The above implication of BMBU regarding the history effects on decision uncertainty (as summarized in Figure 3G,H) was grounded on our model simulation of the trial-to-trial states of the latent variable *u*_*t*_. And this simulation was carried out using the model best fit to the participants’ choice data.

Thus, the validity of the simulated history effects critically relies on how accurately our model fitting procedure recovers the true states of *u*_*t*_ when the model complexity (e.g., number of parameters) and the experimental design (e.g., number of trials) are taken into account. To address this issue, we conducted a variable recovery test (see STAR methods for details) and confirmed that the recovered states of *u*_*t*_ well matched their true states (*R*^2^ = 0.91 ± 0.0038; mean ± 95% confidence interval).

As stressed earlier, BMBU’s ability to exhibit the pre-congruence effect relies on the assumption that the memory recall of a stimulus becomes increasingly noisier as trials elapse. To examine the necessity of this assumption in accounting for the pre-congruence effect, we created three model variants in which the memory decay process is modified such that the boundary is inferred based on either only the immediately preceding stimulus (*β*_*i*_ = 0 for *i* > 1 in Equation 2), all the previous stimuli but with equal weights (*κ* = 0 in Equation 1), or the future stimuli instead of the previous stimulus (*r*_*t+i*_ is substituted for *r*_*t-i*_ in Equation 2). These model variants were all poorer than the original BMBU in predicting the trial-to-trial choice variability (Figure S1A). More importantly, they all failed to display neither the repulsive bias nor the pre-congruence effect on decision uncertainty (Figure S1B,C), which suggests that the assumption of memory decay is required for BMBU to account for the pre-congruence effect.

### Pre-congruence effect in RT

To verify the pre-congruence effect predicted by BMBU, we first probed RT, a behavioral measure widely accepted as a correlate of decision uncertainty^16,18,24^, by regressing it concurrently onto the pre-congruence (*S*_*t*-1_ ∗ *C*_*t*_), current-congruence (*S*_*t*_ ∗ *C*_*t*_), and choice-repetition (*C*_*t*-1_ ∗ *C*_*t*_) factors (Figure 4). Participants’ RTs increased as the current choice became congruent with the previous stimulus, in support of the pre-congruence effect 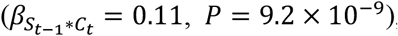, and decreased as the current choice became congruent with the current stimulus, in support of the current-congruence effect (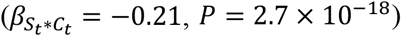, *P* = 2.7 × 10^−18^) (Figure 4A,B). By contrast, the RTs were unaffected by whether the previous choice was repeated in the current trial (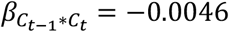, *P* = 0.81; Figure 4B). Notably, the size of the pre-congruence effect on the RTs was substantial and tantamount to 31% of the current-congruence effect (Figure 4B). The pre-congruence effect could be traced back to three trials (Figure 4C), consistent with the history effect on choice—the repulsive bias (Figure 2C).

**Figure 4.**
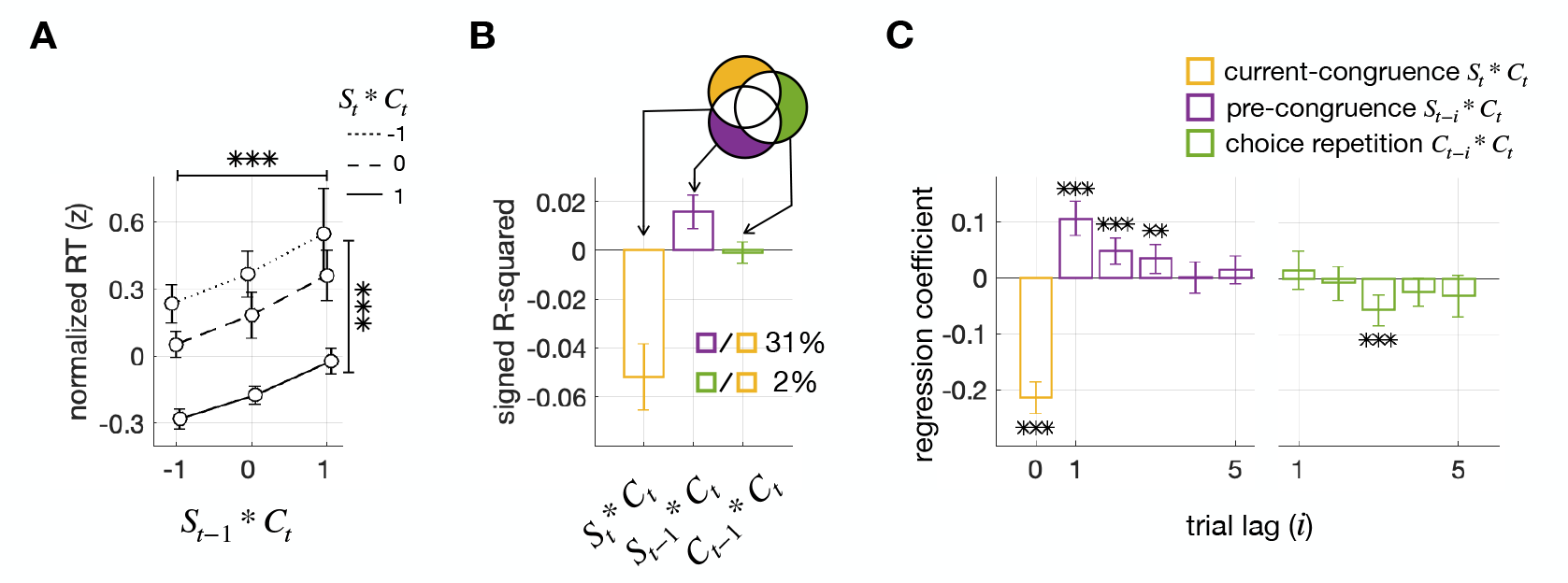
Pre-congruence and current-congruence effects on RT. (A) Changes in RT as a function of the pre-congruence and current-congruence factors. Normalized measures of RT are plotted against the levels of the congruence between the current choice and the immediately preceding stimulus (*S*_*t*-1_ ∗ *C*_*t*_), separately for each level of the congruence between the current choice and the current stimulus (*S*_*t*_ ∗ *C*_*t*_). The asterisks on top and side indicate the statistical significance (*P* < 0.001) of the pre-congruence (*S*_*t*-1_ ∗ *C*_*t*_) and current-congruence (*S*_*t*_ ∗ *C*_*t*_) effects, respectively. Their significance was assessed by the multiple regressions of RT onto the two congruence factors (*S*_*t*-1_ ∗ *C*_*t*_ and *S*_*t*_ ∗ *C*_*t*_) and the choice-repetition factor (*C*_*t*-1_ ∗ *C*_*t*_). (B) Explained variance analysis. Bars plot the RT variance uniquely explained by the current-congruence factor (*S*_*t*_ ∗ *C*_*t*_; yellow), the pre-congruence factor (*S*_*t*-1_ ∗ *C*_*t*_; purple), and the choice-repetition factor (*C*_*t*-1_ ∗ *C*_*t*_; green), respectively, as illustrated by the Venn diagram. The percentage scores quantify the ratio of the variances explained by the pre-congruence and choice-repetition factors, respectively, to the variance explained by the current-congruence factor. (C) Multiple regression analysis. Bars plot the regression coefficients for the current-congruence factor ((*S*_*t*_ ∗ *C*_*t*_; yellow), the pre-congruence factor with trial lag *i* (*S*_*t*-*i*_ ∗ *C*_*t*_; purple), and the choice-repetition factor with trial lag *i* (*C*_*t*-*i*_ ∗ *C*_*t*_; green). The asterisks indicate the significance of regression coefficients: ∗, *P* < 0.05; ∗∗, *P* < 0.01; ∗∗∗, *P* < 0.001.

### Relationship between the pre-congruence effect and previously reported history effects involving decision confidence

Next, having confirmed the presence of the pre-congruence effect in RT, we considered the possibility that the pre-congruence effect is associated with, or can even be explained away by, previously reported history effects that involve decision uncertainty. Two such history effects are an autocorrelation in decision confidence between consecutive trials^50^ and a tendency to repeat choices when decision confidence is high^17,18^. We indeed confirmed these two effects in our dataset: (i) the RT in the current trial (*RT*_*t*_) increased as the RT in the previous trial (*RT*_*t*-1_) increased (Figure 5A 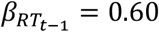, *P* = 3.0 × 10^−4^); (ii) the probability of repeating choices (*C*_*t*-1_ = *C*_*t*_) increased as the RT in the previous trial (*RT*_*t*-1_) decreased and vice versa (Figure 5B; *t* = 3.14, *P* = 0.0032), which means that the repetitive and the alternative choice biases are facilitated after fast and slow RT in the preceding trial, respectively. If the facilitated choices involved faster *RT*_*t*_, *RT*_*t*_ would be faster as choice is repetitive after fast *RT*_*t*-1_, whereas *RT*_*t*_ would be faster as choice is alternative after slow *RT*_*t*-1_. This modulatory effect of the choice history on the association between *RT*_*t*_ and *RT*_*t*-1_ is observed by regressing *RT*_*t*_ concurrently onto *RT*_*t*-1_ and the interaction term *C*_*t*-1_ ∗ *C*_*t*_ × *RT*_*t*-1_ (Figure 5C; 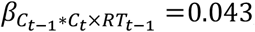, *P* = 0.0046; 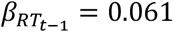, *P* = 2.1 × 10^−4^). Therefore, we examined whether the two history effects of RT identified in consistent with previous studies can explain away the pre-congruence effect.

**Figure 5.**
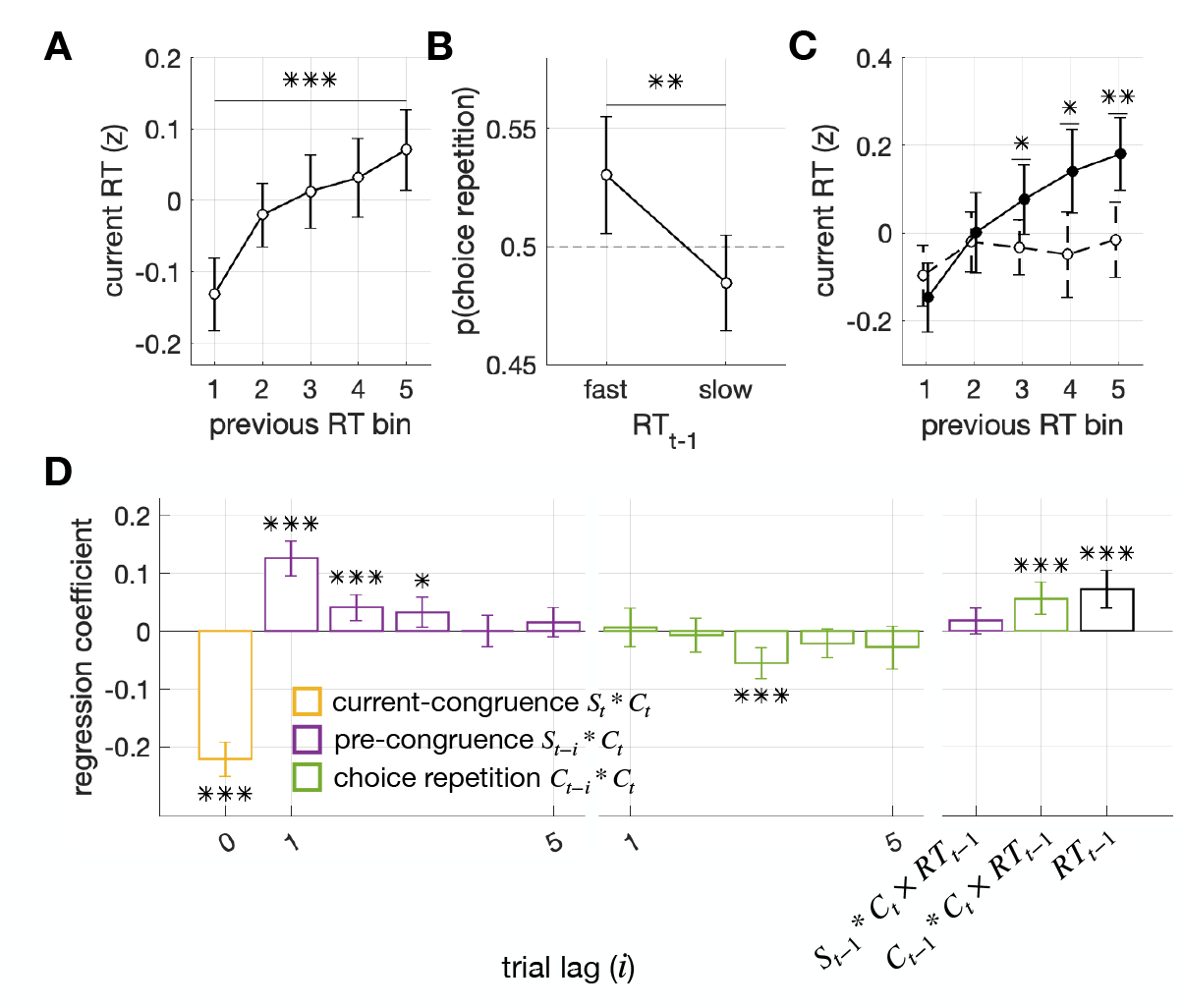
Co-existence of pre-congruence and current-congruence effects with other confidence-related history effects. (A) Positive correlation in RT between consecutive trials. The RT in the current trial was plotted against the binned RTs in the previous trial. The asterisks indicate the significance (*P* < 0.001) of the linear regression of the current RT (*RT*_*t*_) onto the previous RT (*R*_*t*-1_). (B) Effect of previous RT on choice repetition. The probability of repeating the previous choice in the current trial is plotted against the split-half levels of the previous RT. The asterisks indicate the significant difference in choice repetition between these two levels (*P* < 0.01, two-sided pair-wise t-test). (C) Modulatory effects of choice repetition on the relationship between previous and current RTs. The RT in the current trial was plotted against the binned RTs in the previous trial, separately when consecutive choices were repeated (*C*_*t*-1_ ∗ *C*_*t*_ = 1; solid circles with solid line) and when alternated (*C*_*t*-1_ ∗ *C*_*t*_ = −1; empty circles with dashed line). The asterisks indicate the significant differences in current RT between the choice repetition and alternation conditions at given bins of previous RT (two-sided pair-wise t-tests; ∗, *P* < 0.05; ∗∗, *P* < 0.01). (D) Multiple regression analysis with additional regressors. The multiple regression analysis depicted in Figure 4C was repeated but with three more regressors: the RT in the previous trial (*RT*_*t*-1_) and the two interaction terms, one reflecting the modulation of the previous RT by the pre-congruence factor ((*S*_*t*-1_ ∗ *C*_*t*_) × *RT*_*t*-1_) and the other reflecting the modulation of previous RT by the choice-repetition factor ((*C*_*t*-*i*_ ∗ *C*_*t*_) × *RT*_*t*-*i*_). The asterisks indicate the significance of regression coefficients: ∗, *P* < 0.05; ∗∗, *P* < 0.01; ∗∗∗, *P* < 0.001.

The two effects require us to check whether the influence of the pre-congruence (*S*_*t*-1_ ∗ *C*_*t*_) on the current RT remains significant when the previous RT (*RT*_*t*-1_) and the modulatory effects of the previous RT (*C*_*t*-1_ ∗ *C*_*t*_ × *RT*_*t*-1_) are taken into account. To meet these two requirements, we conducted the previous regression analysis on *RT*_*t*_ while including three additional regressors: the previous RT (*RT*_*t*-1_), its modulation of choice repetition (*C*_*t*-1_ ∗ *C*_*t*_ × *RT*_*t*-1_), and its modulation of pre-congruence (*S*_*t*-1_ ∗ *C*_*t*_ × *RT*_*t*-1_) (Figure 5D). We found that the current RT was significantly regressed onto the previous RT (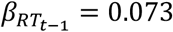, *P* = 4.9 × 10^−5^) and its modulation of choice repetition (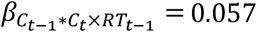, *P* = 2.3 × 10^−4^), consistent with the two effects mentioned above, but not onto the previous RT’s modulation of the pre-congruence (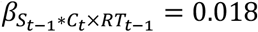, *P* = 0.13). More importantly, as in the original regression, where those three additional regressors were not included, the influence of the pre-congruence on the current RT remained significant even after three trials elapsed (compare Figure 5C to Figure 4C). These results indicate that the two previously reported history effects involving decision uncertainty do not explain away but co-exist with the pre-congruence effect.

### Robustness of the pre-congruence effect to sensory modality, stimulus granularity, inter-trial interval, and feedback presence

According to BMBU, the pre-congruence effect is to occur under various experimental conditions as long as the boundary for binary classification needs to be inferred from previously seen stimuli. To test this prediction, we analyzed three more datasets in our laboratory, which differ from the main dataset in sensory modality, stimulus granularity, inter-trial interval, or feedback presence (see Table 1 for specifications). Despite these differences, all these auxiliary datasets displayed the pre-congruence effect that is qualitatively similar to that in the main dataset for three important aspects (Figure 6). First, the current RT increased as the current choice became increasingly more congruent with the previous stimulus (Figure 6A1-3). Second, the pre-congruence effect was substantial in size, explaining 10% to 24% of the variance in RT that was explained by the current-congruence effect (Figure 6B1-3). Lastly, except for the dataset where trial-to-trial feedback was used, the pre-congruence effect remained significant even when trials elapsed by as far as five (Figure 6C1-3). These results suggest that the pre-congruence effect is a robust phenomenon, at least being generalizable to the experimental conditions used for our auxiliary datasets.

**Table 1.** Task protocols for the five datasets used in the current study.

| dataset name<br>(number of subjects) |  | stimulus | stimulus number | trial-to-trial<br>feedback | inter-trial<br>interval (s) |
| --- | --- | --- | --- | --- | --- |
| main dataset | fMRI (18) | ring | 3 | x | 13.2 |
|  | eye tracking (23) | ring | 3 | x | 13.2 |
| fine-grained pitch (58) |  | pitch | continuous | x | 2 |
| fine-grained ring (58) |  | ring | continuous | x | 2 |
| feedback (30) |  | ring | 5 | o | 2.5 |

**Figure 6.**
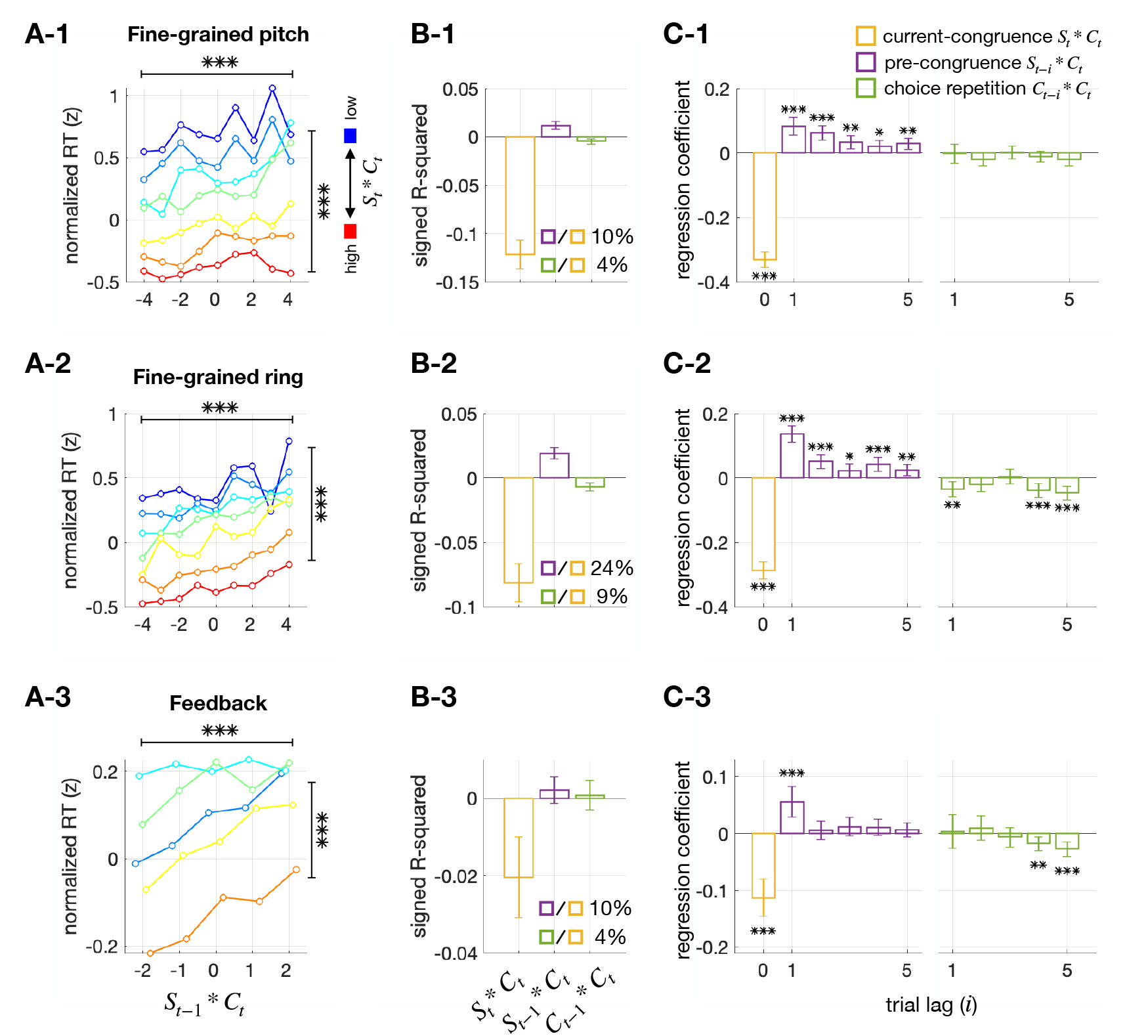
Pre-congruence and current-congruence effects on RT in three other datasets acquired with different task protocols. The analyses shown in Figure 4 were repeated on three additional datasets. While these datasets used the same binary classification task as the main dataset, they varied in several task protocols, including stimulus feature, stimulus granularity, inter-trial interval, and presence of trial-to-trial feedback. The name and details of each dataset are specified in Table 1. The summarized results of the datasets named ‘fine-grained pitch,’ ‘fine-grained ring,’ and ‘feedback’ in Table 1 are presented in the top (1), middle (2), and bottom (3) rows of panels. For datasets ‘fine-grained pitch’ and ‘fine-grained ring,’ where continuous stimuli were used, the stimuli were discretized into nine bins for illustrative purposes. In doing so, the data points at the two lowest bins defined for the current-congruence factor (*S*_*t*_ ∗ *C*_*t*_) are not shown because they were the trials where incorrect choices were made despite strong stimulus evidence and thus very rare. (A) Changes in RT as a function of the pre-congruence and current-congruence factors. The format is identical to that of Figure 4A. (B) Explained variance analysis. The format is identical to that of Figure 4B. (C) Multiple regression analysis. The format is identical to that of Figure 4C.

### Pre-congruence effect in the fMRI activity of the dorsal anterior cingulate cortex

Previous studies have identified neural correlates of decision uncertainty in various non-human animal^14,49,51-53^ and human^15,35-44^ brain regions. Since we acquired fMRI measurements from eighteen of the participants while they performed the ring size classification task, we searched their brains for neural correlates of decision uncertainty and examined whether those neural correlates display the pre-congruence effects as BMBU implies.

We began by regressing the fMRI data solely onto the simulated trial-to-trial states of decision uncertainty produced by BMBU (*u*_*t*_). As a result, we identified its neural correlates in the dorsal anterior cingulate cortex (dACC) and insula (Figure 7A; Table S1). In both regions, the fMRI responses showed the highest correlation with *u*_*t*_ at 5.5 s after the stimulus onset (Figure S2, panels in the top and middle rows), which makes senses, given the hemodynamic delay in fMRI signal, because decision uncertainty is expected to be greatest at the moment of decision-making. Thus, as we did for the RT data, we regressed the trial-to-trial fMRI responses at that time point onto the three types of regressors (Figure 7B-D). The regression onto the current-congruence factor was significant in both regions (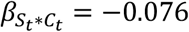, *P* = 0.0016 for the dACC; 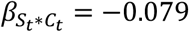, *P* = 1.4 × 10^−5^ for the insula). By contrast, the regression onto the pre-congruence factor was significant only in the dACC (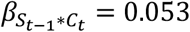, *P* = 0.0037 for the dACC; 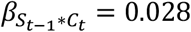, *P* = 0.083 for the insula). The pre-congruence effect in the dACC was 46% of the size of the current-congruence effect (Figure 7C-1) and could be traced back to two trials (Figure 7D-1), similar to the RT findings. Furthermore, by expanding the regression analysis for the remaining within-trial time points, we confirmed that the pre-congruence effect is most pronounced at the moment of decision-making (Figure 7E-1).

**Figure 7.**
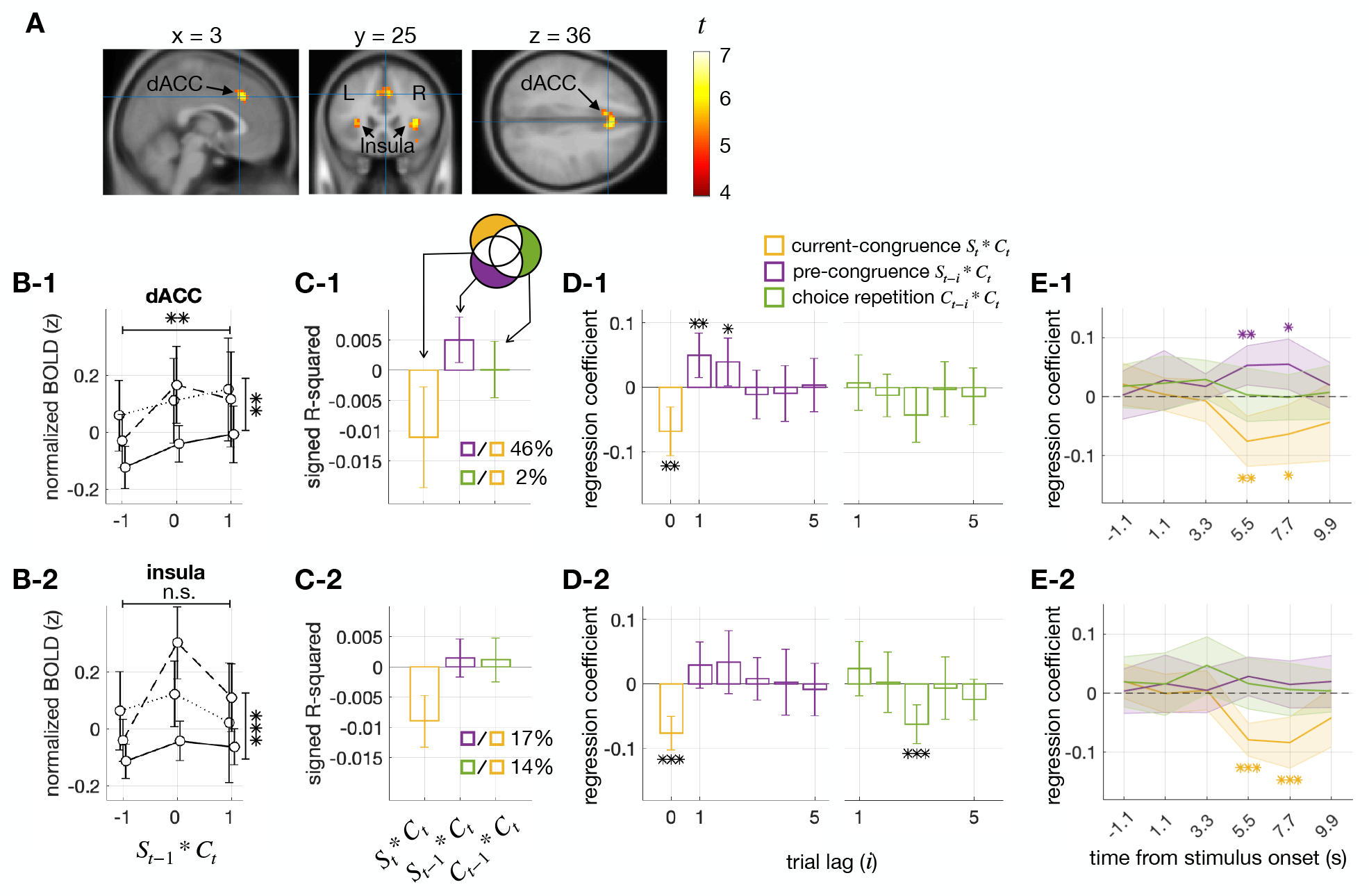
Pre-congruence and current-congruence effects on cortical activity. (A) Definition of the cortical regions of interest (ROIs). The ROIs were defined by carrying out a two-sided Student’s t-test with a false discovery rate of 0.05 and identifying voxel clusters with a minimum number of 15 contiguous voxels whose BOLD responses significantly predicted the model estimates of decision uncertainty. The *t*-statistics of these clusters are overlaid on the sagittal (left), coronal (middle), and axial (right) slices of the anatomical image of a template brain. The panels below (B-E) present the results for the ROIs in the dACC and insula in the top (1) and bottom (2) rows, respectively. (B) Changes in BOLD responses as a function of the pre-congruence and current-congruence factors. The format is identical to that of Figure 4A. Note that both congruence effects were significant in the dACC (1), whereas only the current-congruence effect was significant in the insula (2). (C) Explained variance analysis. The format is identical to that of Figure 4B. Note that the ratio of the variances explained by the pre-congruence to the variance explained by the current-congruence factor was more substantial in the dACC (1) than in the insula (2). (D) Multiple regression analysis. The format is identical to that of Figure 4C. Note that the regression coefficients of the pre-congruence factor were significant in the dACC (1) but not in the insula. (E) Time courses of the current-congruence, pre-congruence, and choice-repetition effects on cortical activity. The regression coefficients of the current-congruence (yellow; *S*_*t*_ ∗ *C*_*t*_), pre-congruence (purple; *S*_*t*-1_ ∗ *C*_*t*_), and choice-repetition (green; *C*_*t*-1_ ∗ *C*_*t*_) factors are plotted over the time frames relative to the stimulus onset. Note that the emergence of the significant pre-congruence effect is aligned with the time frames associated with decision-making only in the dACC not in the insula. See also Figure S2,3 and Table S1.

### Pre-congruence effect in pupillary responses

Previous studies have identified physiological correlates of decision uncertainty in pupillary responses^18,25-34^: the higher the decision uncertainty becomes, the larger the pupil grows in size. For the remaining twenty-three participants from whom fMRI data were not acquired, we tracked their pupillary responses during the task. Thus, we repeated the regression analysis on the pupil-size data to further verify the pre-congruence effect.

Consistent with the fMRI responses, the pupillary responses also showed the highest correlation with *u*_*t*_ at the moment of decision-making (Figure S2, panels in the bottom rows). Thus, as we did for the fMRI data, we regressed the trial-to-trial pupillary responses at that time point onto the three types of regressors (Figure 8A-C). The regressions onto the current-congruence and pre-congurence factors were both significant (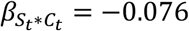, *P* = 0.0016; 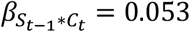, *P* = 0.0037). The pre-congruence effect in pupil-size was 35% of the size of the current-congruence effect (Figure 8B) and could be traced back to one trial (Figure 8C). Similar to the fMRI data, we also confirmed that the pre-congruence effect in pupil-size is most pronounced at the moment of decision-making (Figure 8D).

**Figure 8.**
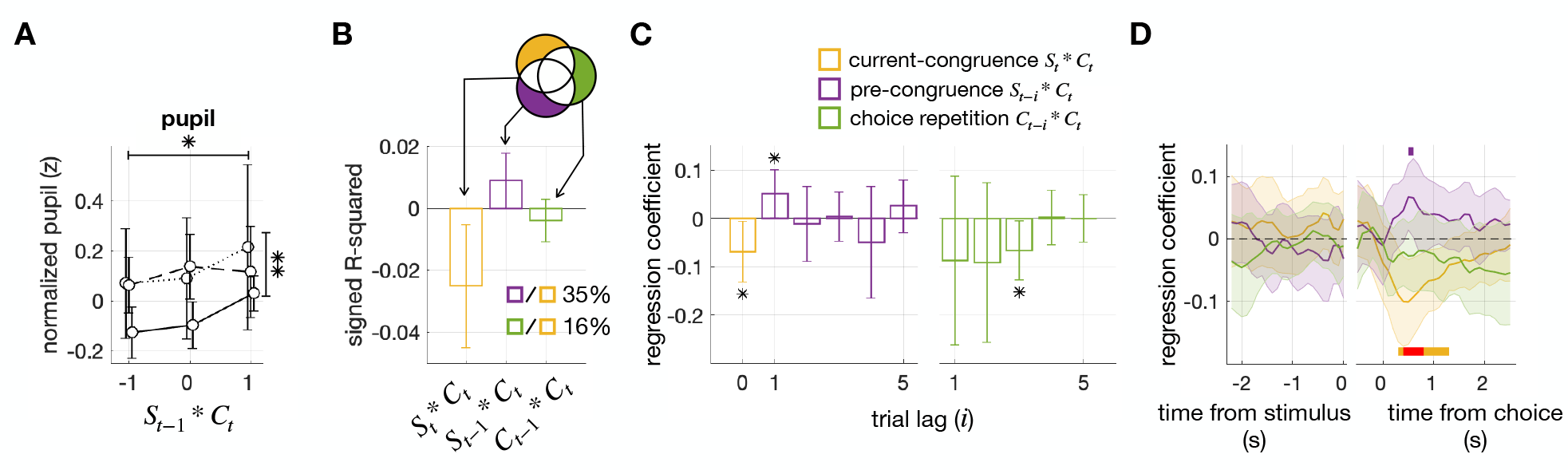
Pre-congruence and current-congruence effects on pupil size. (A) Changes in pupil size as a function of the pre-congruence and current-congruence factors. The format is identical to that of Figure 4A. (B) Explained variance analysis. The format is identical to that of Figure 4B. (C) Multiple regression analysis. The format is identical to that of Figure 4C. (D) Time courses of the current-congruence, pre-congruence, and choice-repetition effects on pupil size. The regression coefficients of the current-congruence (yellow; *S*_*t*_ ∗ *C*_*t*_), pre-congruence (purple; *S*_*t*-1_ ∗ *C*_*t*_), and choice-repetition (green; *C*_*t*-1_ ∗ *C*_*t*_) factors are plotted over the time frames relative to the stimulus onset. The colored horizontal bars demarcate the time points where the regression coefficients of the pre-congruence and current congruence factors were significant (two-sided Student’s *t*-test; purple, *P* < 0.05 for the pre-congruence factor; yellow, *P* < 0.05 and red, *P* < 0.01 for the current-congruence factor). See also Figure S2.

## DISCUSSION

Binary classification entails setting a boundary that divides quantities of interest into two even halves. Assuming that human brains are bounded by memory decay, BMBU prescribes a Bayesian solution of how the class boundary should be updated from their noisy recalls of recent stimuli. This solution implies that the class boundary is incessantly drawn towards recent stimuli over decision-making trials. As shown in our prior work^12^, this boundary shift toward recent stimuli effectively explains the repulsive bias, a history effect of previous stimuli on current choices. Here, we deduced that the same boundary shift also leads to another, previously unrecognized, history effect: as the current choice becomes congruent with the previous stimulus, the decision uncertainty accompanying the current choice increases, which we refer to as the pre-congruence effect. We confirmed the pre-congruence effect in three well-known correlates of decision uncertainty: RT, dACC activity, and pupil size. Our analysis of the RT data indicates that the pre-congruence effect is not only robust to the changes in sensory modality, stimulus granularity, inter-trial interval, and trial-to-trial feedback but also cannot be attributed to the previously reported history effects associated with decision uncertainty. By concurrently verifying the implications of BMBU on the repulsive bias and pre-congruence effect, our work points to boundary updating as a common source of the history effects both on choice and decision uncertainty.

### Novelty and uniqueness of the pre-congruence effect

Previous studies have shown that decision uncertainty, or its behavioral correlate such as RT, is influenced by various history factors. However, these factors differ from the pre-congruence factor, an extent to which the previous stimulus is congruent with the current choice (*S*_*t*-1_ ∗ *C*_*t*_). Below, we will identify these other history factors respectively and explain why they cannot explain away the pre-congruence effect.

Rahnev et al. (2015)^50^ reported that subjective confidence reports are positively correlated across consecutive trials, dubbed ‘confidence leak.’ Here, the history factor is the subjective confidence report in the previous trial and thus includes neither the previous stimulus nor the current choice, unlike the pre-congruence factor. We showed that the current RT (*RT*_*t*_) is regressed concurrently onto the previous RT (*RT*_*t*-1_), a proxy of the previous confidence factor, and the pre-congruence factor (Figure 5A,D), which suggests the independence between the confidence leak and the pre-congruence effect.

Donner and his colleagues^17,18^ reported that the tendency to repeat the previous choice in the current trial increases as the decision uncertainty accompanying the previous choice decreases. Here, decision uncertainty is not a target of the history effect but a history factor—a modulatory factor modifying the influence of the previous choice (*C*_*t*-1_) on the current choice (*C*_*t*_), to be exact.

Nevertheless, the association between choice repetition (*C*_*t*-1_ \**C*_*t*_) and decision uncertainty somehow might have contributed to the pre-congruence effect, which bears on the relationship between decision uncertainty and the pre-congruence (*S*_*t*-1_ \**C*_*t*_), via the correlation between the previous stimulus (*S*_*t*-1_) and choice (*C*_*t*-1_). However, this possibility seems unlikely because the pre-congruence effect remained almost unaffected by the previous RT’s modulatory effect (Figure 5D).

There are two other history effects on RT where the previous stimuli or their relationship with the current stimulus exert their influence on RT. First, patterns in preceding stimulus sequences, such as repetition and alternation, have been reported to affect the current RT^54^. The history factor is defined by the second-order relationship among past stimuli in this phenomenon, whereas the pre-congruence effect is defined by the respective relationships of past stimuli with the current choice. Second, RT is known to decrease as the current stimulus becomes similar to the previous one, dubbed ‘priming effect’^55,56^. The history factor is the congruence between the previous and current stimuli in the priming effect, whereas the history factor of pre-congruence effect is the congruence between the previous stimuli and the current choice. Even when considering the correlation between the current stimulus and choice, the bias in the priming effect is opposite to that in the pre-congruence effect: RT decreases in the priming effect but increases in the pre-congruence effect as a function of the congruence. Thus, the pre-congruence effect is distinct from these two history effects of stimuli on RT.

In a phenomenon called ‘post-error slowing,’ RT tends to increase following an error in the previous trial^57^. Here, the history factor is trial-to-trial feedback. However, we showed that the pre-congruence effect robustly occurred with and without trial-to-trial feedback (Figure 6).

Lastly, the pre-congruence effect should not be confused with a phenomenon where RT tends to be faster when the previous choice is repeated^58^. Here, the history factor is ‘choice repetition’—i.e., the congruence between the previous and current choices (*C*_*t*-1_ \**C*_*t*_). The regression of RT onto this choice repetition factor was also present in our datasets but much less substantial than the pre-congruence factor (Figures 4B,C and 6B,C). We suspect that these weak effects of choice repetition could have been caused by the counteract of the positive effect of motor repetition on RT^59^ against the negative effect of choice repetition^58^ because choices and motor responses were confounded in our experiment.

Grounded on the comparisons above, we conclude that the pre-congruence effect is a novel history effect that cannot be explained away by any previously reported history effects on decision uncertainty or RT.

### Boundary updating, a principled and integrative account of the history effects on choice and decision uncertainty

Perceptual decision-making involves committing to a proposition about a particular world state. This commitment is expressed as a categorical choice and is accompanied by a continuous level of decision uncertainty. From a normative perspective, both ‘choice’ and ‘decision uncertainty’ are rooted in a common source, ‘decision variable.’ The choice is determined by which option is more supported by the decision variable, while the decision uncertainty is determined by how the chosen option is more supported by the decision variable than the unchosen option^23,48,49^. This close relationship between choice and decision uncertainty implies that the history effect on choice systematically relates to the history effect on decision uncertainty.

However, previous studies have rarely inspected these two types of history effects together, instead mainly focusing on the history effect on choice^8,13,17,60-65^. Although a few studies have explored the relationship between the history effects on choice and RT in a descriptive manner^58,66^, they have not provided any clear explanation for these relationships. Our work offers a principled account by pointing to “boundary updating towards recent stimuli” as a common source of the history effects on choice and decision uncertainty: previous stimuli concurrently induce the repulsive bias in choice and the pre-congruence effect on decision uncertainty by attracting the class boundary towards themselves (Figure 1).

It is worth noting the ‘normative’ nature of updating the class boundary towards recent stimuli, given memory decay in the brain. If the brain’s memory recalls of past stimuli were perfect, then its knowledge about stimulus distribution would stabilize quickly after a reasonable number of trials, resulting in a boundary that is not easily influenced by recent stimuli. In such a situation, the pre-congruence effect is considered ‘irrational’ because any ‘rational’ decision-maker should only rely on current sensory input. However, due to memory decay, the brain has to rely on its memory recalls of more recent, thus more reliable, stimuli to better infer the actual stimulus distribution, resulting in a boundary that is incessantly drawn to recent stimuli. In this regard, the pre-congruence effect can be considered a by-product of the memory-bounded brain’s ‘normative’ solution to infer the actual stimulus distribution from noisy memory recalls of recent stimuli. Looking at the pre-congruence effect from this perspective offers a fresh understanding of how history effects on decision uncertainty can be reinterpreted as rational decision-making. For instance, the presence of history effects on confidence reports has been considered an indication of decision-makers’ metacognition inefficiency^67^. However, if we consider the normative nature of boundary updating, the absence, not the presence, of the history effect on confidence reports may indicate their metacognitive inefficiency.

Lastly, it is important to note that the pre-congruent effect is not only statistically significant but also substantial in size. In fact, it ranged from 10% to 46% of the current-congruency effect in our datasets. This implies that if boundary updating is not incorporated into the estimation of decision uncertainty, the trial-to-trial estimates of decision uncertainty substantially deviate from their actual states. Consistent with this implication, cortical activity in any single voxel did not significantly correlate with decision uncertainty when a model variant lacking the boundary updating process was used to define decision uncertainty (Figure S3). Additionally, incorporating boundary updating into the definition of decision uncertainty made our model estimates of decision uncertainty sensitive enough to identify the dACC, but not the insula, as a genuine neural correlate of decision uncertainty based on the presence of the pre-congruence effect (Figure 7). Given that previous efforts to identify the neural correlate of task difficulty have found it difficult to distinguish between these two areas^11,16,68,69^, considering the pre-congruence effect is expected to facilitate the search for the ‘genuine’ neural substrate of decision uncertainty.

### Limitations of the Study

It is important not to confuse the ‘class boundary’ with the ‘decision boundary’ in signal detection theory^70,71^. The class boundary represents the center of a stimulus distribution, while the decision boundary is the optimal boundary value that maximizes the objective function of any given decision-making task. Therefore, unlike the class boundary, the decision boundary can be adjusted not only by previous stimuli^72^ but also by the value and base rate of choices^73,74^. This means that decision uncertainty can be influenced by non-stimulus history factors as well. In this regard, BMBU should be considered an account of the history effects that arise from previously experienced stimuli. More work needs to be done to comprehensively understand the history effects that come from diverse sources.

In addition, the formalism of BMBU has a restriction of only being suitable for binary classification on quantities that vary along a single dimension. Therefore, it would be worth incorporating the decision boundary into BMBU because the formalism of signal detection theory is proficient in dealing with quantities defined in more than one dimension^70^.

Our results can be said to address the history effect on decision confidence because the statistical definition of decision confidence (*f*_*t*_) is the probability that the current choice is correct^23,48,49^ which is the reciprocal relationship to decision uncertainty (*u*_*t*_ = 1 – *f*_*t*_). However, it is hard to argue that our work addressed the history effect on the decision confidence that is subjectively defined, which is often measured by the ratings provided by decision-makers^48^. This is because the metacognitive ability may interfere with the performance of the confidence rating in this self-report paradigm^67^. Further research is needed to investigate whether the pre-congruence effect is present in self-rated decision confidence.

## ACKNOWLEDGEMENTS

This research was supported by the Research Grant from Seoul National University (339-20220013), the Brain Research Program (No. NRF-2021R1F1A1052020, NRF-2017M3C7A1047860), and the Basic Research Laboratory Program (NRF-2018R1A4A1025891) and the Basic Science Research Program (RS-2023-00276729) through the National Research Foundation of Korea (NRF) funded by the Ministry of Science and ICT.

## AUTHOR CONTRIBUTIONS

Conceptualization, H.L. and S.-H.L.; Methodology, H.L. and S.-H.L.; Investigation, H.L. and S.-H.L.; Writing – Original Draft, H.L.; Writing – Review & Editing, H.L. and S.-H.L.; Funding Acquisition, H.L. and S.-H.L.; Resources, S.-H.L.; Supervision, S.-H.L.

## DECLARATION OF INTERESTS

The authors declare no competing interests.

### Declaration of Generative AI and AI-assisted technologies in the writing process

During the initial drafting process, H.L. utilized ChatGTP to verify the naturalness of his phrases and sentences. Afterward, the corresponding author, S.-H.L., thoroughly revised the initial draft and made the necessary edits to the content. S.-H.L. takes full responsibility for the publication’s content.

## STAR METHODS

### KEY RESOURCE TABLE

| REAGENT or RESOURCE | SOURCE | IDENTIFIER |
| --- | --- | --- |
| Deposited Data |  |  |
| Raw and analyzed data | This paper | https://github.com/Heeseung-Lee/iScience-2023.git |
| Software and algorithms |  |  |
| MATLAB | MathWorks | https://www.mathworks.com/ |

### RESOURCE AVAILABILITY

#### Lead contact

Further information and requests for resources should be directed to and will be fulfilled by the lead contact Sang-Hun Lee.

#### Materials availability

Besides data and MATLAB codes, this study did not generate any new reagents or materials.

#### Data and code availability

- All data have been deposited at GitHub and are publicly available. The DOI is listed in the key resources table.
- All original code has been deposited at GitHub and is publicly available. The DOI is listed in the key resource table.
- Any additional information required to reanalyze the data reported in this paper is available from the lead contact upon request.

### EXPERIMENTAL MODEL AND STUDY PARTICIPANT DETAILS

We utilized three datasets used for previous papers of our group^12,75-77^ and two new datasets using fine-grained size and pitch stimuli. The fMRI dataset^12^ consisted of 18 participants (9 females) aged 20-30 years. The eye tracking dataset^75,76^ comprised 23 participants (11 females) aged 18-36 years. The trial-to-trial feedback dataset^77^ included 30 participants (13 females) aged 18-30 years. The two new datasets using fine-grained stimuli shared the same subjects which included 58 (38 females) aged 18-34 years. The experimental procedures were approved by the Research Ethics Committee of Seoul National University, and all participants provided informed consent and were unaware of the study’s objectives.

### METHOD DETAILS

#### Task procedure

The main data set used the following task procedure (Figure 2A): Participants were instructed to fixate at the center of the screen and classify a brief (0.3s) ring-shaped stimulus as ‘small’ or ‘large’ within 1.5s after stimulus onset by pressing the left or right key, respectively. The timing and identity of each key press were recorded. Trials were separated by 13.2s, and participants were given feedback on their performance after each run of 26 trials.

Before the main runs, participants completed 54 practice trials followed by 180 threshold-calibration trials. During the threshold-calibration trials, which were separated by 2.7s, participants received trial-to-trial feedback based on the boundary with a radius of 2.84°. A Weibull function was fit to the psychometric curves obtained from the threshold-calibration trials using a maximum-likelihood procedure. The size threshold *Δ*° (i.e., the size difference between medium ring and large or small ring) associated with a 70.7% correct proportion was estimated from the fitted Weibull function. The mean and standard deviation of the estimated size threshold were 0.023 and 0.0078, respectively. One of three estimated ring sizes (2.84 − *Δ*°, 2.84°, 2.84 + *Δ*°) was shown on each trial. Participants had extensive training on the task before participating in the main experiments.

Two of the three supplementary datasets were collected from 58 participants, who performed pitch and ring-size classification tasks on separate sessions, respectively, with 315 trials using fine-grained stimuli randomly sampled from normal distributions with an inter-trial interval of 2s and without trial-to-trial feedback for each task. The other dataset was collected from 30 participants, who performed the same classification task over 5 daily sessions with 1,700 trials using ring stimuli of 5 discrete sizes (3.84°, 3.92°, 4.00°, 4.08°, 4.16°), with trial-to-trial feedback and an inter-trial interval of 2.5s.

#### A Bayesian model for boundary updating (BMBU)

##### The generative model

The generative model is the observers’ causal account for noisy sensory measurements. Specifically, the true size of the ring, *S*, is the cause of a noisy measurement, *m*_*t*_, on the current trial. This measurement becomes noisier as more trials pass, resulting in a noisier measurement of the value of *S* on trial *t* − *i*, which is denoted as *r*_*t*-*i*)_ (Figure 3A). The generative model can be defined using three probabilistic terms: the prior probability of *S, P*(*S*), the likelihood of *S* given *m*_*t, P*(*m*_*t*_|*S*), and the likelihood of *S* given *r*_*t*-*i*_, *P*(*r*_*t*-*i*_|*S*). All three terms are modeled as normal distribution functions, which are characterized by mean and standard deviation parameters, *μ* and σ. Specifically, the prior is characterized by *μ*_0_ and σ_0_, the likelihood for *m*_*t*_ is characterized by 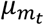 and 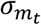, and the likelihood for *r*_*t*-*i*_ is characterized by 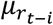 and 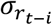. The mean parameters of the likelihood functions are identical to the observed sensory measurements, *m*_*t*_ and *r*_*t*-*i*_. Therefore, the four parameters that need to be learned are *μ*_0_, σ_0_, 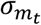, and 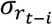. The parameter 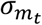 is assumed to be constant across different values of *m*_*t*_ and across trials and is therefore reduced to a constant value σ_*m*_. The parameter 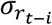 is assumed to increase with the number of elapsed trials, and it is modeled as a parametric function of *κ*, where *k* > 0: 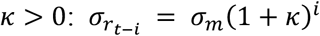. As a result, the generative model is fully specified by the four parameters, which are represented as Θ = [*μ*_0_, σ_0_, σ_*m*_, *k*].

##### Stimulus inference (s)

To estimate the value of *S* on a current trial based on the sensory measurement *m*_*t*_, a Bayesian approach was employed, where the mean of the posterior distribution of *S* given *m*_*t*_, denoted as *P*(*S*|*m*_*t*_), was considered as the Bayesian estimate *s*_*t*_. This posterior distribution is a conjugate normal distribution of the prior and the likelihood of *S* given the sensory evidence *m*_*t*_. The Bayesian estimate *s*_*t*_ and its standard deviation 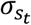 were calculated as follows:

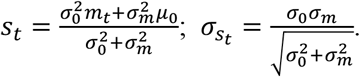

##### Class boundary inference (b)

The Bayesian observer estimates the value of the class boundary on a current trial, *b*_*t*_, by taking the mean of the posterior function based on a given set of retrieved sensory measurements, 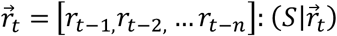, The maximum number of measurements that can be retrieved, *n*, is set to 7 because it is much longer than the effective trial lags of the previous stimulus effect (Figure 2). The posterior function, 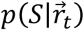, is a conjugate normal distribution of the prior and likelihoods of *S* given the evidence 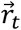:

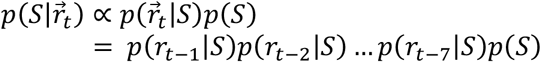

whose mean and standard deviation were calculated based on the knowledge of how the retrieved stimulus becomes noisier as trials elapse:

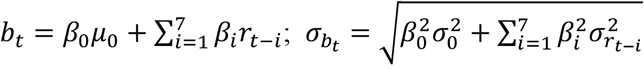

where 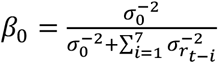 and 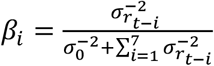.

##### Deduction of choice (c) and decision uncertainty (u)

For each trial, if *s*_*t*_ exceeds *b*_*t*_, the simulated choice *c*_*t*_ is recorded as ‘large’ (or 1); otherwise, it is recorded as ‘small’ (or -1). In addition, we defined the decision uncertainty, *u*_*t*_, which represents the likelihood that the current decision will be incorrect. Specifically, *u*_*t*_ is calculated as follows:

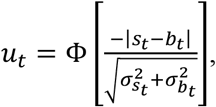

where Φ stands for the standard normal cumulative function.

#### A constant-boundary model

The constant-boundary model comprises two parameters, the bias of the class boundary, *μ*_0_, and the measurement noise, σ_*m*_. In this model, we assumed that stimulus estimates, *s*_*t*_, were sampled from a normal distribution, *𝒩*(*S*_*t*_, σ_*m*_), where *S*_*t*_ represents the physical size of the stimulus. The uncertainty in each stimulus sample was assumed to be equal to σ_*m*_, that is, 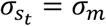. Furthermore, we assumed that the boundary estimate, *b*_*t*_, was a constant value, *μ*_0_, with 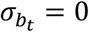.

#### BOLD signal

MRI data were collected using a 3 Tesla Siemens Tim Trio scanner equipped with a 12-channel Head Matrix coil at the Seoul National University Brain Imaging Center. Stimuli were generated using MATLAB (MathWorks) in conjunction with MGL (http://justingardner.net/mgl) on a Macintosh computer. Observers looked through an angled mirror attached to the head coil to view stimuli displayed via an LCD projector (Canon XEED SX60) onto a back-projection screen at the end of the magnet bore at a viewing distance of 87cm, yielding a field of view of 22×17°.

For each observer, we acquired three types of MRI images, as follows: (1) 3D, T1-weighted, whole-brain images (MPRAGE; resolution, 1×1×1mm; field of view (FOV), 256mm; repetition time (TR), 1.9s; time for inversion, 700ms; time to echo (TE), 2.36; and flip angle (FA), 9°), (2) 2D, T1-weighted, in-plane images (MPRAGE; resolution, 1.08×1.08×3.3mm; TR, 1.5s; T1, 700ms; TE, 2.79ms; and FA, 9°), and (3) 2D, T2*-weighted, functional images (gradient EPI; TR, 2.2s; TE, 40ms; FA, 73º; FOV, 208mm; image matrix, 90×90; slice thickness, 3mm with 10% space gap; slice, 32 oblique transfers slices; bandwidth, 790Hz/ps; and effective voxel size, 3.25×3.25×3.3mm).

The images of individual observers were normalized to the MNI template using the following steps: motion correction, coregistration to whole-brain anatomical images via the in-plane images^78^, spike elimination, slice timing correction, normalization using the SPM DARTEL Toolbox^79^ to 3×3×3mm voxel size, and smoothing with 8×8×8mm full-width half-maximum Gaussian kernel. All the procedures were implemented with SPM8 and SPM12 (http://www.fil.ion.ucl.ac.uk.spm)^80,81^, except for spike elimination for which we used the AFNI toolbox^82^. The first six frames of each functional scan (the first trial of each run) were discarded to allow hemodynamic responses to reach a steady state. Then, the normalized BOLD time series at each voxel, each run, and each subject were preprocessed using linear detrending and high-pass filtering (132s cut-off frequency with a Butterworth filter), conversion into percent-change signals, correction for non-neural nuisance signals by regressing out the mean BOLD activity of the cerebrospinal fluid (CSF), and standardization by subtracting the mean activity and by dividing the standard deviation.

To define the anatomical mask of CSF, probability tissue maps for individual participants were generated from T1-weighted images, normalized to the MNI space, and smoothed as were done for the functional images by using SPM12, and then averaged across participants. Finally, the location of CSF was defined as respective groups of voxels in which the probability was more than 0.5.

Unfortunately, in a few of the sessions, functional images did not cover the entire brain. We investigated a brain region only when it does not loss any session. As a result, some voxels in the temporal pole, ventral orbitofrontal, and posterior cerebellum were excluded from data analysis.

#### Pupillometry

Stimuli were presented in a dimly lit room on a gamma-linearized 22-inch CRT monitor (Totoku CV921X CRT monitor) operating at vertical refresh rate of 180Hz and a spatial resolution of 800×600 pixels. Stimuli were generated using MATLAB (MathWorks) in conjunction with MGL (http://justingardner.net/mgl) on a Macintosh computer. Observers viewed the monitor at a distance of 90cm while their binocular eye positions were sampled at 500Hz by an infrared eye tracker (EyeLink 1000 Desktop Mount, SR Research; instrument noise, 0.01° RMS). The LED illuminator and camera were positioned side by side, at a distance of 65cm from the observer, and angled toward the observer’s face to ensure that infrared light illuminated both eyes and was being reflected from both eyes and imaged on the camera sensor.

When measuring gaze position using video-based methods, eye blinks can interfere with the accuracy of the data, as pupil and gaze information are not available during these periods. We identified eye blinks based on three criteria: (i) missing pupil data for either eye, (ii) pupil-size measurements with unrealistically large fluctuations (>50 units per sample), or (iii) substantial deviation (>20°) of gaze position from the screen center. Data collected immediately before and after an eye blink (±200 ms) were likely contaminated and therefore we linearly interpolated the pupil-sizes and the three gaze positions (*x, y*, and 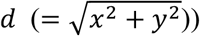 before and after the blink events.

To minimize confounding effects from gaze position on pupil-size measurements, we processed the blink-free samples of pupil-size and three gaze positions time series by linear detrending, band-pass filtering (0.01Hz to 4Hz cut-off frequency with a Butterworth filter), and resampling to 10Hz. The resampled time series of pupil-size was then orthogonalized from the three resampled gaze positions up to their fourth powers (total 12 regressors) to extract any confounding in the pupil-size originating from the gaze position^75^. The orthogonalized pupil-size time series was then standardized by subtracting the mean and dividing the standard deviation of the time series. The pupil-sizes of the eyes were averaged, and trials were epoched and baseline corrected by subtracting the baseline pupil-size averaged across 0ms to 500ms from the stimulus onset. We chose this baseline time window because we assumed that the decision-related pupil signal would be initiated between -200ms to 300ms from the stimulus onset and because the latency of the empirical responses was around 200ms^83^. Finally, the baseline corrected pupil-sizes were aligned by the stimulus onset or the choice following previous studies^18,84^.

### QUANTIFICATION AND STATISTICAL ANALYSIS

#### Model fitting

To estimate the parameters of the generative model for each human participant, we used maximum likelihood estimation. Specifically, we sought to find the values of Θ = [*μ*_0_, σ_0_, σ_*m*_, *k*] that maximized the sum of log-likelihoods across *T* individual choices made by the human observer, 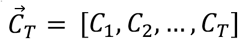:

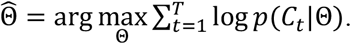

For each participant, we estimated the parameters of the generative model in the following steps. Initially, we used a MATLAB function, fminseachbnd, to find local minima of parameters with the iterative evaluation number set to 50. To acquire a set of candidate parameter estimates, we repeated this step by choosing 1,000 different initial parameter sets randomly sampled within uniform prior bounds. In the second step, we selected the top 20 candidate sets based on goodness-of-fit (sum of log-likelihoods) and searched for the minima using each of those 20 sets as initial parameters. We increased the iterative evaluation number to 100,000 and set tolerances of function and parameters to 10^−7^ for reliable estimation. Using the parameters fitted via the second step, we repeated the second step once more. Then, we selected the parameter set that showed the largest sum of likelihoods as the final parameter estimates. We excluded the first trial of each run and trials with reaction times less than 0.3 seconds from further analyses. This exclusion is necessary because the first trial of each run does not have its previous trial, which is necessary for investigating the repulsive influence. Additionally, a response made during the stimulus’s initial presentation (0∼0.3s) may be too hasty to reflect a normal cognitive decision-making process.

#### Definition of congruence

The term ‘congruence’ refers to the product of the variable of interest and the current choice, where choices were represented as 1 for ‘large’ and -1 for ‘small’ and stimuli were represented as 1, 0, and -1 for large, medium, and small sizes, respectively. For instance, if the current choice was ‘large (1)’ and the previous stimulus was a ‘small’ size (−1), then the congruence would be -1 (1 * -1). In the supplementary data where there were five stimuli, the stimuli were encoded as [-2, -1, 0, 1, 2].

#### Estimation of decision uncertainty

By fitting the model parameters 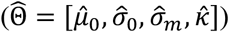 separately for each human participant, we were able to create a Bayesian observer tailored to each individual. By conducting the experiment again with these Bayesian observers using the same stimulus sequences presented to their human counterparts, we obtained a sufficient number (10^6^ repetitions) of simulated choices, *c*_*t*_, and decision uncertainty values, *u*_*t*_. These values were determined based on the corresponding number of stimulus estimates, *s*_*t*_, and boundary estimates, *b*_*t*_, for each Bayesian observer. Finally, we took the averages across those 10^6^ simulations as the final outcomes. When estimating *u*_*t*_ for the observed choice *C*_*t*_, we only included simulation outcomes where the simulated choice *c*_*t*_ matched the observed choice *C*_*t*_.

#### Variable recovery test

To ascertain the validity of our procedure of estimating the trial-to-trial states of decision uncertainty *u*_*t*_, we ran simulations to check how accurately the true states of *u*_*t*_ are recovered by the procedure. This variable recovery test is crucial because the accuracy of the model estimates of *u*_*t*_ is a prerequisite for correctly identifying the brain regions where BOLD responses correlate with decision uncertainty and the time points when pupil size measurements correlate with decision uncertainty. The variable recovery test was carried out with the following aim and procedure.

The variable recovery test aimed to evaluate the correspondence between the *true* and *recovered* states of *u*_*t*_ on a trial-to-trial basis for each of the individual observers, who are likely to vary in parameters. To achieve this aim, it is important to generate a comprehensive set of *true* trial-to-trial states of *u*_*t*_ using a realistic set of model parameters that reflect the range of parameters of actual observers. To do so, we first created 256 different sets of parameter values by taking the possible combinations of the four different values of each of the four model parameters (*μ*_0_, σ_0_, σ_*m*_, *k*), where the four different values corresponded to the 20, 40, 60, and 80 percentiles of the parameter values fitted to the choices of the observers (N=41) who participated in either the fMRI (N=18) or the eye-tracking (N=23) experiments. Second, we acquired the synthetic choices and the *true* states of decision uncertainty (*u*_*t*_) by plugging one of the 256 parameter sets into BMBU and simulating it on the actual stimulus sequence used for one of the 41 observers. Third, we fitted the 4 parameters of BMBU to those synthetic choices for each observer.

Fourth, we simulated the *recovered* states of *u*_*t*_ using those fitted model parameters on the identical stimulus sequence used for generating the *true* states of *u*_*t*_. Fifth, we calculated the R-squared between the *true* and *recovered u*_*t*_s to assess how reliably our model fitting procedure can recover the *true* states of the model variables. Sixth, we repeated the second to fifth steps for all the remained 255 parameter sets and used the R-squared averaged across all the 256 parameter sets as the performance measure of the recovery test for the one observer of interest. Finally, we repeated the second to sixth procedures for all the remaining 40 observers and used the average R-squared and its confidence interval across all the 41 observers as the performance measure of the recovery test.

#### Definition of the signed R-squared

The signed R-squared is a measure that takes into account the direction of the relationship between a variable of interest and other variables in a multiple regression model. It is calculated by multiplying the sign of the regression coefficient with the uniquely explained variance of the variable. The uniquely explained variance is obtained through variance partitioning analysis^85,86^, which involves comparing the R-squared values of the full regression model that includes all variables and the reduced regression model that excludes the variable of interest. For instance, to calculate the signed R-squared of variable *x*, we first calculate the R-squared of the full model 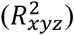 that includes variables *x, y*, and *z*. Next, we compute the R-squared of the reduced model 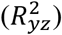 that includes variables *y* and *z* only. The uniquely explained variance of 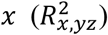 is obtained by subtracting 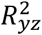 from 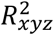. Finally, the signed R-squared of *x* is defined as 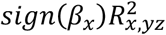, where *sign*(*β*_*x*_) is the sign of the regression coefficient *β*_*x*_ of the multiple regression model with variables *x, y*, and *z* as regressors.

#### Searching for the brain signals correlated with u_t_

To explore the brain regions that showed a correlation between BOLD signals and *u*_*t*_, we conducted a regression analysis for each participant and voxel. The response variable for the regression model was the preprocessed BOLD signals concatenated across runs. We used three explanatory vectors in the regression model. The first vector was created by convolving the canonical hemodynamic function with *u*_*t*_ for each trial. The second vector was created by convolving the canonical hemodynamic function with a constant value of 1, and both vectors were standardized. The third vector contained only a single value of 1. The first regression coefficient indicated how well *u*_*t*_ predicted the BOLD signal for each voxel, and by calculating this coefficient for every voxel, we generated a map of regression coefficients for each participant and the entire brain.

To determine whether the average coefficient of *u*_*t*_ across participants significant, we performed a two-sided Student *t*-test on each voxel. We then corrected the *P* values of the entire brain for false discovery rate (FDR) ^87^ (Figure S3). The total number of voxels in the entire brain was 90481, and we defined the brain areas significantly correlated with *u*_*t*_ as the voxel clusters covering a region larger than 15 contiguous voxels and having FDR-corrected *P* values less than 0.05. For the ROI analysis, we averaged the preprocessed BOLD signals across individual voxels within an ROI.

## SUPPLEMENTAL INFORMATION

**Figure S1.**
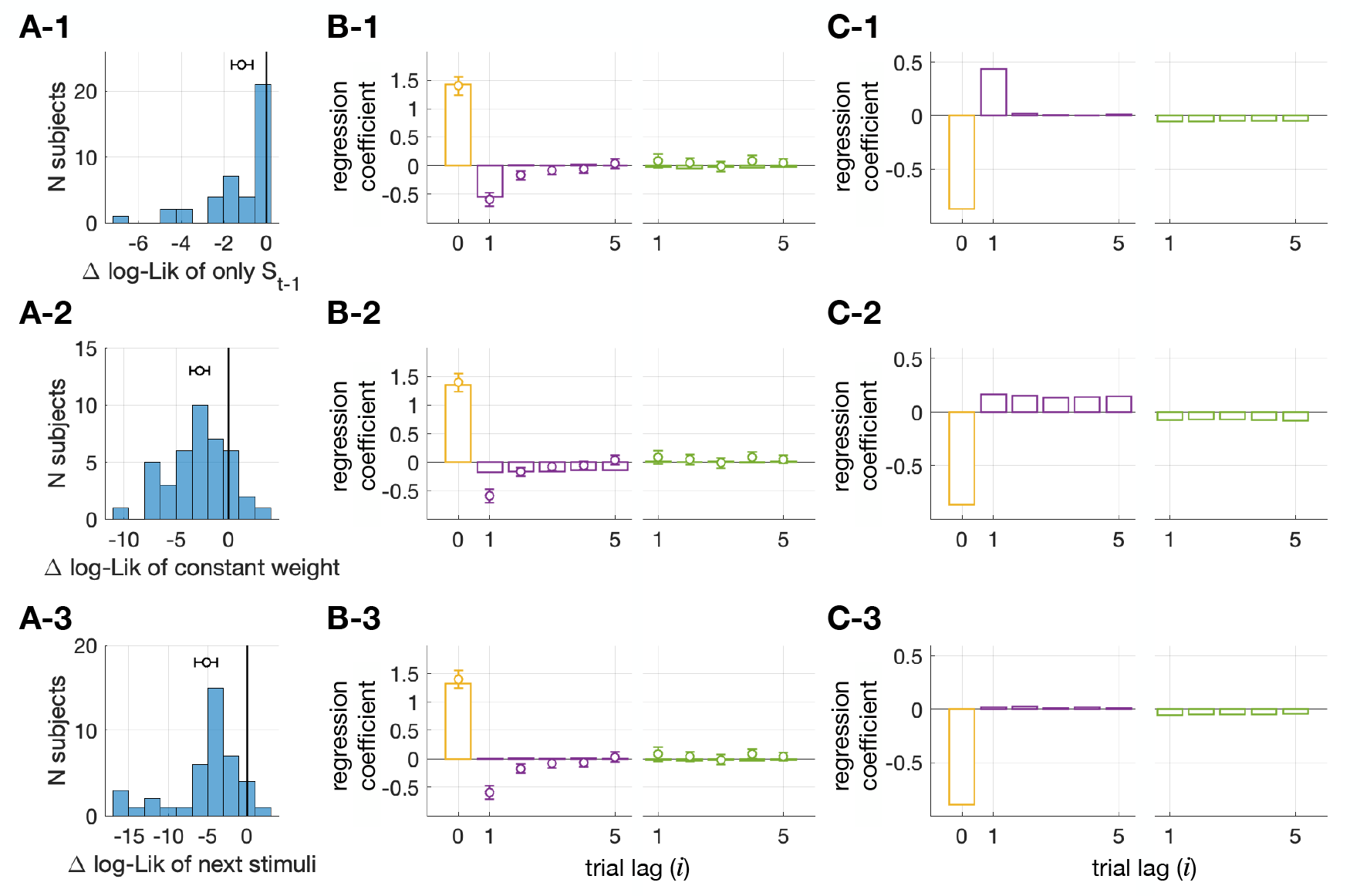
Necessity of the ‘memory-decay’ assumption in explaining the pre-congruence effect, related to Figure 3. To evaluate the necessity of the assumption that the memory recall of a stimulus becomes increasingly noisier as trials elapse (Equations 1 and 2) in accounting for the pre-congruence effect, we created three variants of the original model (BMBU) by modifying the process of boundary inference, as follows. In the first model variant called ‘limited memory access,’ the memory recall was assumed to be accessible only for the immediately preceding stimulus (*β*_*i*_ = 0 for *i* > 1 in Equation 2). In the second model variant called ‘no memory decay,’ the noisiness of memory recall was assumed not to increase after one trial lag 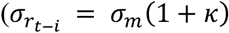 in Equation 1). In the third model variant called ‘prospective memory recall,’ the boundary inference was based on the future stimuli instead of the past stimuli (*r*_*t*-i_ is replaced with *r*_*t*+i_ in Equation 2). These three model variants were compared with BMBU for goodness-of-fit evaluation, the results of which are summarized in A. The multiple regression analyses depicted in Figure 2C and 3H were repeated using the three model variants, the results of which are summarized in B and C, respectively. The summarized results of the ‘limited memory access,’ ‘no memory decay,’ and ‘prospective memory recall’ model variants are presented in the top (1), middle (2), and bottom (3) rows of panels. (A) Goodness-of-fit analysis. The histograms show the frequency (number of observers) distributions of the differences in log-likelihood between BMBU and the model variants. For each observer, the predictive power (AIC) of BMBU was subtracted from those of the model variant. The circles and horizontal error bars represent the means and their 95% confidence intervals of the log-likelihood differences. The log-likelihoods of the model variants were all significantly smaller than that of BMBU (‘limited memory access,’ *t* = −4.7, (*P* = 2.7 × 10^−5^); no memory decay,’ *t* = −6.2, (*P* = 2.2 × 10^−7^); ‘prospective memory recall,’ *t* = −7.3, (*P* = 7.9 × 10^−9^)). (B) Model simulation of repulsive bias demonstrated in logistic regressions. The coefficients in the multiple logistic regression of the current choice (*C*_*t*_) onto the current and previous stimuli (*S*_*t*-i_) and the previous choice (*C*_*t*-i_) are plotted in circles and bars for humans and the models, respectively. (C) Model simulation of pre-congruence and current-congruence effects in linear regressions. The vertical bars represent the coefficients of the multiple linear regression of the simulated decision uncertainty onto the congruences of current choices with the current stimulus (*S*_*t*_ ∗ *C*_*t*_), the previous stimuli (*S*_*t*-i_ ∗ *C*_*t*_), and the previous choices (*C*_*t*-i_ ∗ *C*_*t*_).

**Figure S2.**
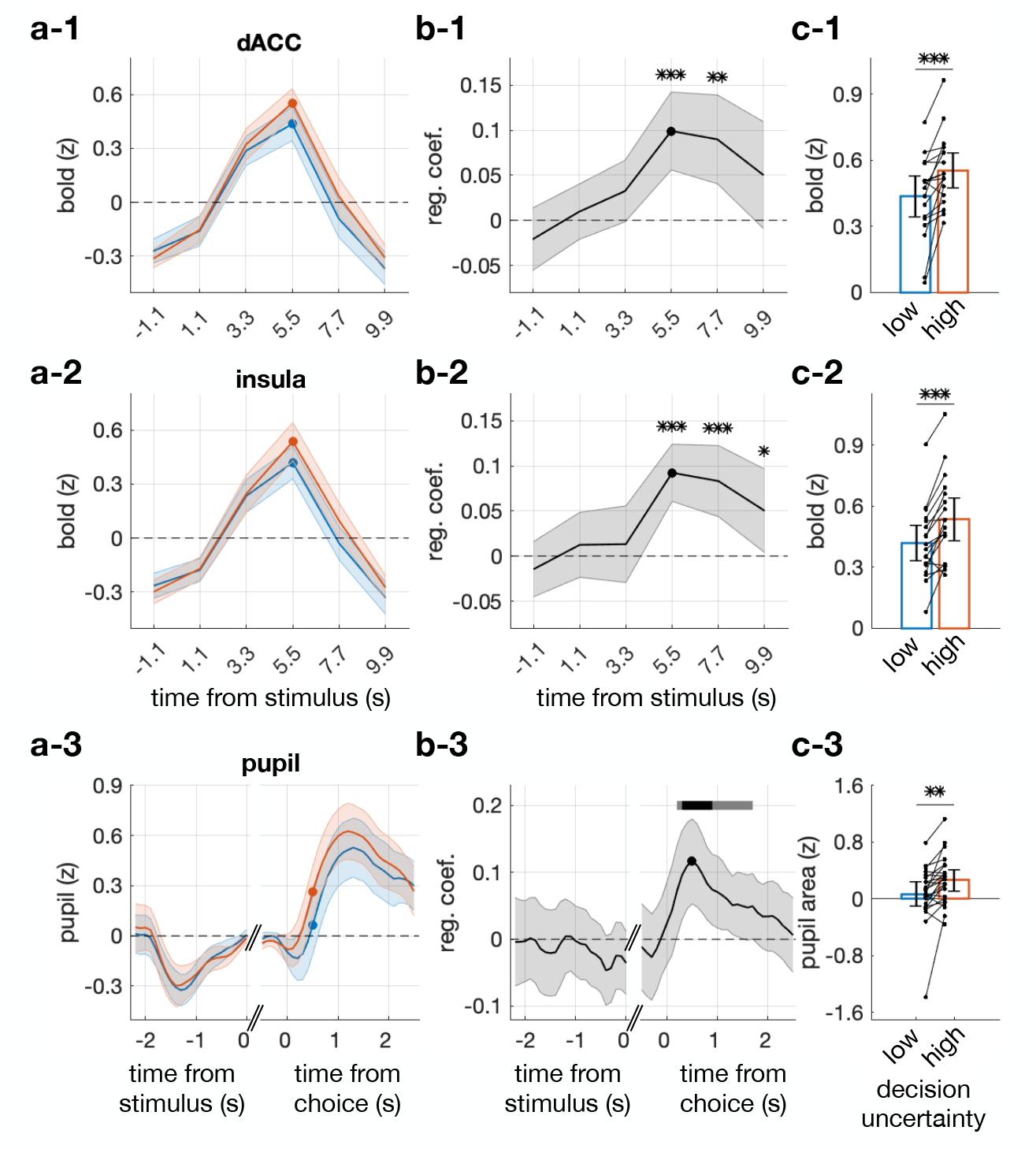
Time courses of the neural and pupillary correlates of the model estimates of decision uncertainty (*u*_*t*_), related to Figure 7,8. To identify the time points where the neural and pupillary correlates of the model estimates of decision uncertainty (*u*_*t*_) are maximally pronounced, we carried out the following analyses on the time series of the BOLD responses in the dACC and insula and of the pupil size measurements. (A) Time courses of BOLD (1,2) and pupil size (3). To visualize the effect of *u*_*t*_ on the BOLD responses and pupil size measurements, we split the trials into the low and high halves in terms of *u*_*t*_. The colors of the symbols, lines, and shades correspond to the two levels of *u*_*t*_ (blue and red for the low and high levels, respectively). (B) Time courses of regression coefficients. We linearly regressed the BOLD responses and pupil size measurements onto *u*_*t*_ in each time point and identified the time points where the significance of those regressions reached their maximum values (the black dots). The coefficients are plotted as a function of time relative to the stimulus onset (1-2) or the time of making choices (3). The *P* values of two-sided Student’s *t*-test of the coefficients are indicated by the horizontal bar (gray: ∗< 0.05; black: ∗∗< 0.01). (C) Differences in BOLD (1,2) and pupil size (3) between the two levels of *u*_*t*_. In the identified time points in B, we compared the signals between trials of low and high halves of *u*_*t*_. The color scheme was identical to that used in A. Each dot pair with a line represents a single observer.

**Figure S3.**
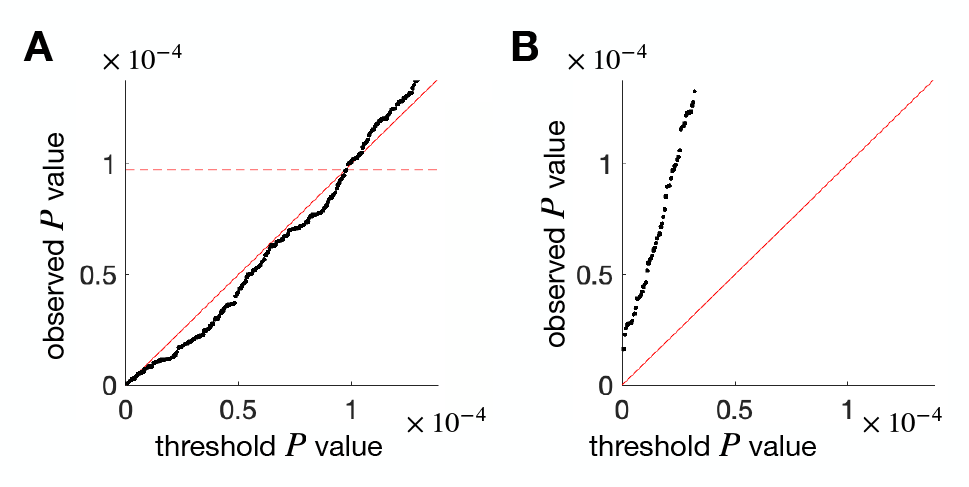
Importance of incorporating the ‘boundary updating’ process into the definition of decision uncertainty in search of its neural correlate, related to Figure 7. To demonstrate how critical the boundary updating process (as depicted in Figure 3A-E) is for the accurate definition of decision uncertainty, we created a constant-boundary model that lacks this process (see STAR Methods for details) and assessed its effectiveness in discovering BOLD correlates of decision uncertainty. Specifically, we regressed the BOLD responses of all brain voxels onto the model estimates of decision uncertainty as defined by BMBU or the constant-boundary model. We then plotted the lowest *P* values of each voxel against the threshold *P* values to control for the false discovery rate (FDR). By doing so, we could determine the significant voxels after controlling for FDR by defining the critical *P* value, which is the maximum *P* value lower than the FDR threshold (indicated by the horizontal dashed line). When BMBU’s estimates of decision uncertainty were used, we found that up to 177 voxels were significant (A), whereas none were significant when the constant-boundary model’s estimates of decision uncertainty were used (B).

**Table S1.** Brain regions where BOLD responses significantly correlate with the model estimates of decision uncertainty (*u*_*t*_), related to Figure 7. To define these regions, we used two criteria: (1) the regression coefficient between the BOLD responses and the model estimates of decision uncertainty was significant (two-sided Student’s *t*-test *P* < 0.05, after controlling for the false discovery rate), and (2) there were at least 15 contiguous voxels that met the first criterion.

| cortical area | continuous voxel (N) | peak voxel |  |  |
| --- | --- | --- | --- | --- |
|  |  | MNI coordinate | *t*-stat | *P*-value |
| dorsal anterior cingulate cortex | 64 | [6, 27, 36] | 6.29 | $8.2 \times 10^{-6}$ |
| left insula | 27 | [−30, 21, 9] | 6.50 | $5.7 \times 10^{-6}$ |
| right insula | 66 | [36, 18, 6] | 7.60 | $7.5 \times 10^{-7}$ |

## Footnotes

1 The class boundary used in this paper is distinct from the ‘decision boundary’ used in signal detection theory or sequential analysis theory. We will explain this distinction in detail in the Discussion section. For the sake of brevity, the class boundary will also be referred to as the ‘boundary.’

